# The functional landscape of patient derived RNF43 mutations predicts Wnt inhibitor sensitivity

**DOI:** 10.1101/2020.03.25.006999

**Authors:** Jia Yu, Permeen Akhtar Bt Mohamed Yuso, Pamela Goh, Nathan Harmston, David M. Epstein, David M. Virshup, Babita Madan

## Abstract

A subset of Wnt-addicted cancers are sensitive to targeted therapies that block Wnt secretion or receptor engagement. RNF43 loss-of-function mutations that increase cell surface Wnt receptor abundance cause sensitivity to Wnt inhibitors. However, it is not clear which of the clinically identified RNF43 mutations affect its function *in vivo*. We assayed 90 missense and 45 truncating RNF43 mutations found in human cancers, using a combination of cell-based reporter assays, genome editing, flow cytometry and immunofluorescence microscopy. Patent-derived xenograft (PDX) models with C-terminal truncating RNF43 mutations were tested for Wnt inhibitor sensitivity. We find that five common germline variants of RNF43 have wild-type activity. The majority of cancer-associated missense mutations in the RING and PA domains are either loss of function or hyperactivating. Hyperactivating mutants appear to function through formation of inactive dimers with endogenous RNF43 and/or ZNRF3. C-terminal truncation mutants including the common G659fs mutant, have discordant behavior in *in vitro* versus *in vivo* assays. PDXs and cell lines with C-terminal truncations show increased cell surface FZD, Wnt/β-catenin signaling and are responsive to PORCN inhibition *in vivo*, providing clear evidence of RNF43 loss of function. In conclusion, RNF43 nonsense and frameshift mutations, including those in the C-terminal domain, and specific missense mutations in RING and PA are loss of function and predict response to upstream Wnt inhibitors in microsatellite stable cancers. This study expands the landscape of actionable RNF43 mutations, potentially extending the benefit of these therapies to additional patients.

**Statement of Significance:** Loss of function RNF43 mutations, first described in pancreatic cancers, drive progression of multiple cancers by increasing cellular sensitivity to Wnt ligands. These cancers are therefore uniquely sensitive to agents such as PORCN inhibitors that block Wnt production. As the PORCN inhibitors and other upstream inhibitors advance into clinical trials it is important to identify the right patients to treat with these upstream Wnt inhibitors. Hence a detailed map of mutations that are actionable is required.

Here we systematically examined a spectrum of 135 patient-derived RNF43 mutations from multiple cancers. Using cell-based reporter assays, genome editing and patient-derived xenografts, we identify rules to guide patient selection. MSS cancers with either truncating mutations anywhere in the gene, including C-terminal truncations around the G659 position, or point mutations in well-defined functional domains, are likely to have RNF43 loss of function and hence a response to therapy.

## Introduction

Wnts are secreted short-range signaling molecules that play key roles during embryonic development and adult homeostasis. As Wnt proteins are synthesized, they are post-translationally palmitoleated by the O-acyltransferase Porcupine (PORCN), a modification that is required for both Wnt secretion and subsequent binding to receptors including Frizzled (FZD) on the receiving cell surface (1-3). Wnt ligand-receptor interaction causes the accumulation of β-catenin in the cytoplasm and its subsequent translocation to the nucleus where it binds to promoters with TCF/LEF to activate target genes.

Mutations that activate the Wnt/β-catenin signaling pathway are found in many cancers. The most common mutations in Wnt signaling affect downstream regulators of β-catenin degradation such as the adenomatous polyposis coli (APC) protein. However, there is a distinct class of cancers that are driven by mutations in genes including *RNF43, ZNRF3* and *RSPO3* (4-9). These cancers have hypersensitivity to Wnt ligands and multiple approaches to treat these Wnt-addicted cancers including PORCN inhibitors, anti-FZD antibodies and anti-R-spondin (RSPO) antibodies have shown efficacy in preclinical studies (10-14).

RNF43 and its paralog ZNRF3 are integral membrane E3 ubiquitin ligases that ubiquitylate cytoplasmic sites on FZD, driving its lysosomal degradation and hence negatively regulating its abundance at the cell surface (7,15). The activity of RNF43 and ZNRF3 at the cell surface is in turn tightly regulated by RSPO ligands and their membrane co-receptors LGR4/5/6 (16,17). The heterotrimeric complex of LGR-RSPO-RNF43/ZNRF3 inhibits ubiquitylation of FZD, thus increasing FZD cell surface abundance and cellular sensitivity to Wnts. This is clinically relevant since loss of function mutations in RNF43 and gain of function mutations in RSPO2 and RSPO3 that increase FZD abundance are found in multiple cancer types and cause Wnt addiction (6,18-20).

One of the goals of precision medicine is to identify actionable mutations. Actionable R-spondin chromosome translocations are identifiable by PCR-based assays (9). However, missense and indel mutations in the *RNF43* gene identified by targeted or NextGen sequencing present a challenge. These mutations occur along the entire *RNF43* coding sequence in multiple cancer types including colorectal, pancreatic, ovarian, biliary tract and gastric (5,21-26) and with an elevated frequency at two G•C tracks in cancers with microsatellite instability (MSI) (5,20). While selected mutations have been studied, a broad survey of how these mutations affect the activity of RNF43 is needed (7,21,25). As PORCN inhibitors advance in clinical trials, delineating which *RNF43* mutations drive Wnt addiction and which are simply passengers is essential to selecting cancer patients who might benefit from this treatment.

In this study, we profiled 135 *RNF43* mutations identified in human cancers. We find that missense mutations in well-defined functional domains, as well as frameshift and nonsense mutations in the N-terminal half of RNF43 are largely loss of function. A subset of these are dominant negative, likely due to their ability to form inactive multimers with nonmutant protein. Multiple truncating mutations in the RNF43 C-terminus, including the recurrent G659Vfs, show loss of function *in vivo* despite variable results *in vitro* and may therefore be actionable in MSS cancer patients. This study of the mutational landscape of RNF43 facilitates the selection of patients who may benefit from upstream Wnt pathway inhibitors.

## Results

### RNF43 mutational landscape analysis

We systematically examined the distribution of RNF43 mutations identified in multiple human cancer types using cBioportal (27,28). Truncating (nonsense and frameshift) mutations were enriched in the first half of the protein, up until the end of the ubiquitin ligase RING domain (Figure 1A) (p=1.45×10-8, binomial test). Missense mutations appeared more evenly distributed, so we investigated whether they were enriched or depleted in specific functional domains. RNF43 has an extracellular protease-associated (PA) domain that interacts with R-spondins (29), a single-pass transmembrane domain (TM), and an intracellular RING domain with E3 ubiquitin ligase function. Both the PA and RING domains of RNF43 had significantly more missense mutations than expected by chance (Figure 1B, p < 0.05), suggesting there is positive selection acting on these domains in cancer. There was no evidence of positive selection for missense mutations in the C-terminal domain (CTD) of RNF43 (Figure 1B). A permutation test confirmed that several amino acids in RNF43 had a significantly higher frequency of truncation or missense mutations than expected by chance alone, including three we identified as common germline variants (Figure S1A-B). Only 16% (N=79) of the RNF43 mutations were shared between different cancer types, indicating that each cancer type has unique RNF43 mutations. A handful of mutations were seen in multiple cancers, the most common being G659Vfs*41 (Figure 1C). Given the diverse nature of the mutational landscape, we asked if there were general principles that determined whether an RNF43 mutation was a loss of function.

**Figure 1.**
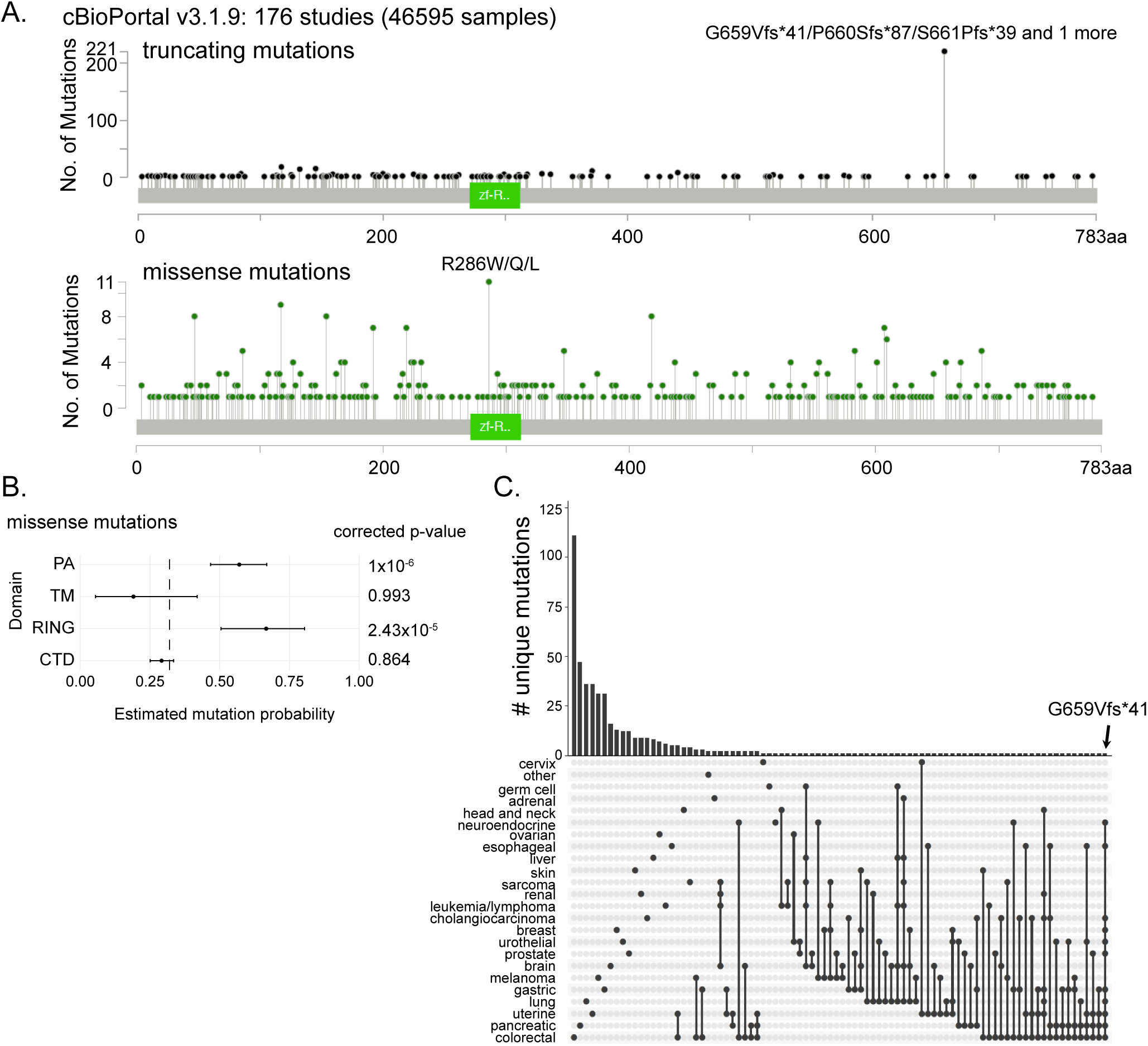
RNF43 mutational landscape. **(A)** *RNF43* truncating (upper panel) and missense mutations (lower panel) reported in cBioPortal human cancer genome database (v3.1.9) from the curated set of non-redundant studies (46,595 samples) show the spread of mutations across the entire gene length with a few hotspots. **(B)** *RNF43* missense mutations are enriched in the PA and RING functional domains (binomial test, Bonferroni corrected p-values). Error bars represent 95% confidence intervals for estimated mutation probability. Dashed line represents the background mutation probability for missense mutations in *RNF43*. **(C)** Upset plot highlights that most *RNF43* mutations (i.e. nonsense, frameshift and missense) are not shared between different cancer types. Single dots indicate the mutations unique to the indicated cancer type, whereas dots with a connecting line indicate the mutations shared between the cancer types. The common G659Vfs*41 mutation is indicated; it is observed in 11 distinct cancers.

### Systematic study of RNF43 missense mutations reveals 30% to be loss-of-function or hyperactivating

It can be difficult to determine *a priori* if a specific missense mutation is a passenger or a driver. To get a better estimate of the actionable RNF43 mutations, we systematically analyzed 90 patient-derived RNF43 missense mutants, selected based on the following criteria. First, we tested the common RNF43 germline variants as these are likely to be found in many tumors. Using the genome aggregation database gnomAD and a cut off of 10% allele frequency we identified five germline variants, with I47V and L418M being the most common with an allele frequency of ∼ 0.4 (Figure 2A) (30). Next, from the cancer-associated missense mutations found in cBioPortal, we focused on mutations that are present in pancreatic cancers, because RNF43 loss of function mutations are well established to render those cancers sensitive to Wnt inhibition (6,19). In addition, we examined a subset of RNF43 mutations present in colorectal cancers with an emphasis on those found in microsatellite stable (MSS) cancers (as assessed by mutational burden) and mutations that are present at high frequencies in other cancer types. We focused on MSS tumors because MSI tumors have multiple additional oncogenic driver mutations and are responsive to alternative therapies such as checkpoint inhibitors (31,32).

**Figure 2.**
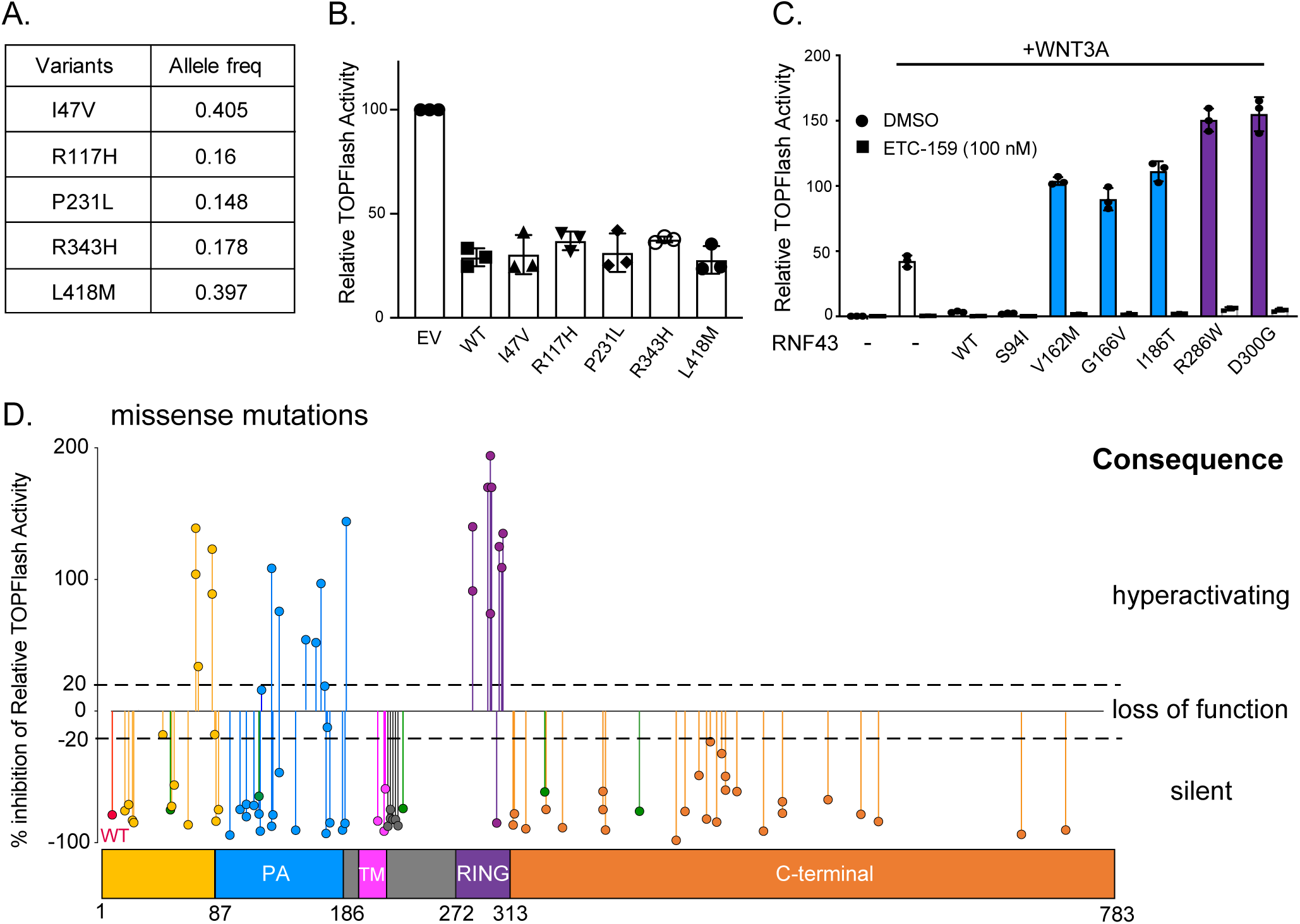
Regulation of Wnt signaling by cancer-associated RNF43 missense mutations. **(A)** Five natural variants of RNF43 with high allele frequency in the general population. **(B)** These germline variants retain wild-type RNF43 activity. HEK293 cells were transfected with Wnt/β-catenin TOPFlash reporter and plasmids expressing WNT3A and the indicated RNF43 variants or empty vector (EV). **(C)** Hyperactivating mutants are sensitive to Wnt inhibition. The activity of RNF43 mutants was assessed as above in B. Cells were treated with 100 nM ETC-159 added 6 hours after transfection where indicated. Data are representative of two independent experiments. **(D)** RNF43 missense mutants in the region around the PA and RING domains are loss of function or hyperactivating. Data represent percentage inhibition of WNT3A induced Wnt/β-catenin reporter activity in HEK293 cells in the presence of the indicated RNF43 mutants. The Wnt/β-catenin reporter activity induced by WNT3A is set as baseline. WT RNF43 inhibits activity by 80-90%,. Activity between −20% to 20% indicates RNF43 loss of function and mutants with activity more than 20% above baseline are hyperactivating. Mutants that inhibit reporter activity more than 20%, are silent mutations. Data are representative of 3 independent experiments. Green dots indicate common germline RNF43 variants.

We used transient expression of RNF43 in a cell-based reporter assay to study the effect of individual *RNF43* mutations. In this assay, wild-type RNF43 reduces Wnt-activated signaling by reducing the abundance of FZD and other Wnt receptors on the cell surface. Benign RNF43 variants should retain activity and reduce Wnt/β-catenin signaling similar to wild-type, while loss-of-function mutants should have no effect. To minimize over-expression artifacts and ensure the sensitivity of the assay, we first titrated the quantity of RNF43 expression plasmid to ensure the amount used is not near saturation (Figure S2A). Patient-derived alterations were introduced into an RNF43 expression vector by site-directed mutagenesis and their ability to regulate WNT induced Wnt/β-catenin reporter activity (TOPFlash) was measured. To assess the robustness of the assay, we compared the activity of selected mutants on signaling stimulated by WNT3A or WNT7B in HEK293 cells, and by WNT3A in a pancreatic cancer cell line, Panc08.13, with WT RNF43 (Figure 2C and Figures S2B-C). Similar results were obtained in all assays, indicating the results obtained are robust and generalizable. Therefore, subsequent screening of RNF43 mutants was performed using HEK293 cells with transfected WNT3A.

Five common RNF43 germline variants were as active as wild-type RNF43 (Figure 2A-B), suggesting they do not contribute to cancer risk and are not actionable. We then assayed the 85 patient-derived missense mutants. The activity of the RNF43 mutants is reported as percent inhibition of Wnt signaling, where wild-type RNF43 produces 80-90% inhibition. The mutants that inhibited signaling less than 20% were defined as loss of function (LOF) and those that actually increased signaling greater than 20% above baseline were termed hyperactivating or dominant negative (Figure 2D). 40% of the missense mutations in the region preceding or in the PA domain (amino acid 87-186) were either loss of function or hyperactivating (Figures 2C-D). Almost all of the mutations in the RING domain (aa 272-313) were hyperactivating. There was also a small cluster of missense mutations in the C-terminus around aa 470-490 that had compromised activity, suggesting an additional functional domain. As the frequency of mutations preceding or in the RING domain are more prevalent, taken together close to 30% of the tested mutations in RNF43 are either loss of function or hyperactivating (Table S1).

In order to test whether these RNF43 hyperactivating mutants are actionable, that is, if they are sensitive to Wnt inhibition, we repeated the reporter assay in the presence of the PORCN inhibitor ETC-159 (19). As expected, PORCN inhibition blocked baseline WNT3A-induced signaling (Figure 2C). Importantly, PORCN inhibition also reduced the enhanced signaling from all tested hyperactivating mutants (V162M, G166V, I186T, R286W and D300G) by >95%. Thus, Wnt secretion is required for the enhanced signaling from RNF43 hyperactivating mutants. Inhibition of Wnt signaling using upstream inhibitors such as anti-FZD antibodies or PORCN inhibitors could therefore be a useful treatment in MSS cancers with either RNF43 loss-of-function or hyperactivating mutations.

### RNF43 regulates the cell-surface abundance of multiple Frizzled proteins

An established molecular function of RNF43 is to ubiquitylate Frizzled (FZD), leading to its endocytosis and lysosomal degradation (7). There are ten FZD genes, and whether RNF43 can act on all of them is not known. Flow cytometry allows assessment of the abundance of FZD at the plasma membrane. The FZDs can be subdivided into four groups based on their sequence similarity (Figure 3A). We first transiently expressed each of the hemagglutinin-epitope (HA)-tagged Frizzled proteins in HEK293 cells. FZD2, 4, 5, 7 and 10, representing three of the four groups, were well expressed and readily detectable in immunoblots with an anti-HA antibody (Figure 3B). To test if RNF43 could regulate the cell surface abundance of these FZDs, they were co-expressed with or without RNF43 in Panc 08.13 cells that have wild-type RNF43 and ZNRF3 and relatively low endogenous FZD levels. As analyzed by flow cytometry (Figure 3C-G), co-expression of wild-type RNF43 reduced the cell surface abundance of each of the five FZDs tested. As an alternative approach to predict whether FZD1, 3, 6, 8 and 9, that were not assessed due to poor expression levels, are also RNF43 targets, we performed multiple sequence alignment of all the ten FZD family members. RNF43 regulates FZD cell surface accumulation by ubiquitination of specific lysine residues (7). The lysines present in both the intracellular loops and the C-terminal domain were largely conserved in all ten FZDs (Figure S2). This suggests that all FZD proteins can be regulated by RNF43.

**Figure 3.**
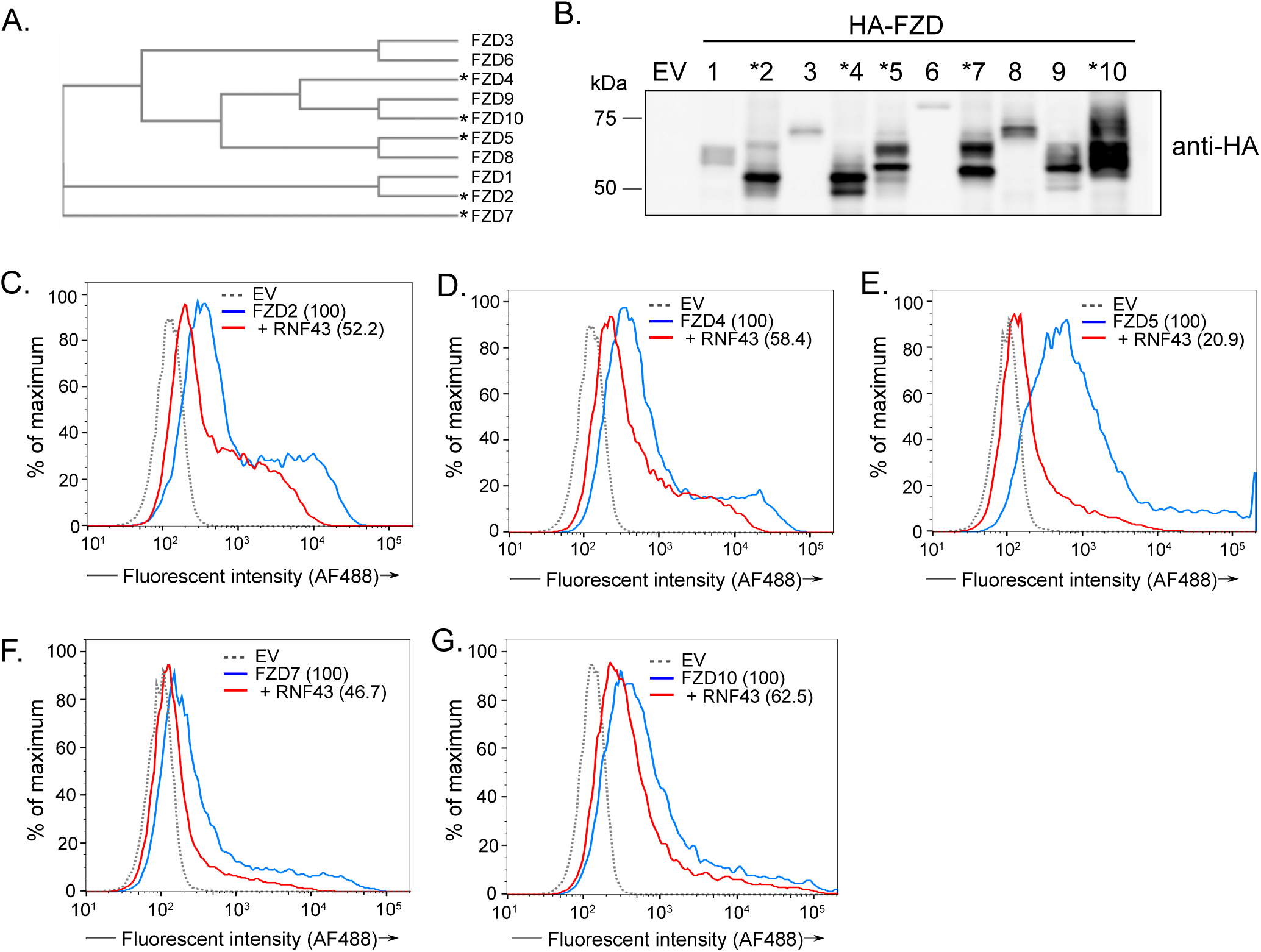
RNF43 regulates cell surface abundance of all tested FZD members. **(A)** Phylogenetic relationship of ten FZD family members. Asterisks indicate the FZD members that were tested by flow cytometric analysis (C-G). **(B)** Expression of FZD family members is variable. Abundance of HA-tagged FZD expressed in HEK293 cells and detected by western blotting with anti-HA antibody. **(C-G)** RNF43 downregulates the cell surface abundance of all tested FZD. The indicated HA-FZDs were expressed in Panc 08.13 cells with or without wild-type RNF43. Cell surface FZD levels were analysed by flow cytometry using Alexa Fluor 488 conjugated anti-HA antibody. Mean fluorescence intensity of cell surface HA-FZD normalized to control is indicated. Data are representative of two independent experiments.

### The activity of RNF43 mutants correlates with their ability to regulate Frizzled abundance, endocytosis and degradation

We used several approaches to test if the activity of RNF43 mutants in regulating Wnt/β-catenin reporter correlated with their ability to regulate endogenous cell surface FZD abundance. First, cell surface FZD levels were measured by flow cytometry using a pan-FZD antibody 18R5 that recognizes five of the ten Frizzled family members (FZD 1, 2, 5, 7 and 8) (11). HEK293 cells have relatively high baseline levels of cell surface FZD (Figure 4A, blue line). Consistent with the ability of RNF43 to reduce Wnt/β-catenin reporter activity, expression of RNF43 significantly reduced the abundance of FZD on the cell surface (Figure 4A, red line). RNF43 PA domain mutant I186T and RING domain mutant R286W that were hyperactivating in reporter assays (Figure 2C) increased the abundance of surface FZD (Figure 4A) while S94I, P118T and A146G mutants that had wild-type activity in Wnt/β-catenin reporter assay behaved like wild-type RNF43 in their ability to reduce FZD abundance (Figure 4B). Similar results were obtained using a different pan-FZD antibody in Panc 08.13 cells (Figure S4A).

**Figure 4.**
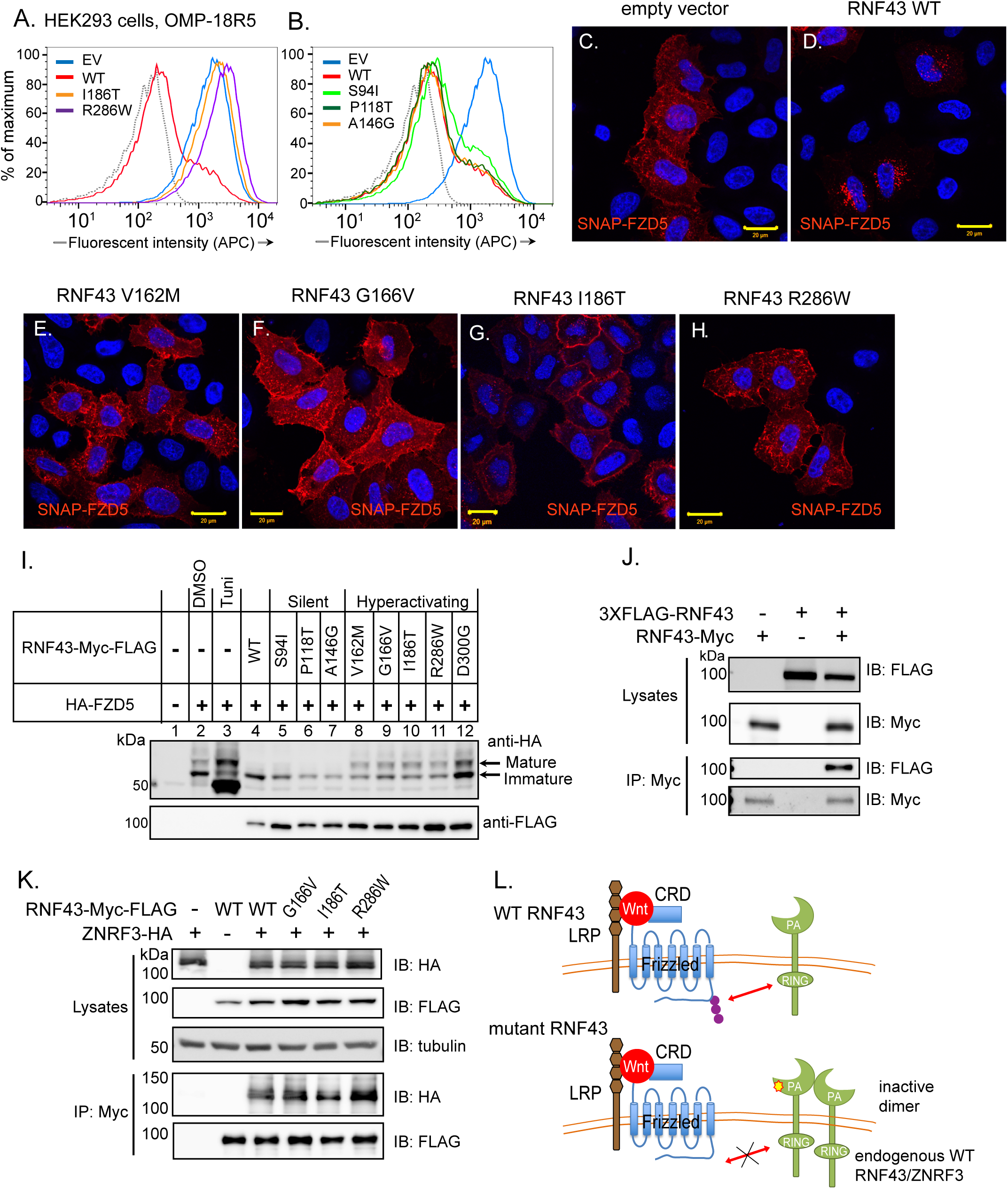
RNF43 hyperactivating mutants form dimers and fail to induce Frizzled endocytosis and degradation. **(A-B)** RNF43 hyperactivating mutants increase cell surface abundance of FZDs. Flow cytometric analysis of endogenous cell surface FZD levels in cells expressing indicated RNF43 mutants. HEK293 cells expressing WT RNF43 or indicated mutants were stained with pan-FZD antibody OMP-18R5. The cells co-expressing GFP were analysed (A) Activating mutants I186T and R286W increase endogenous cells surface FZD abundance. (B) Silent mutants S94I, A146G and P118T downregulate endogenous cell surface FZD comparable to wild-type RNF43. Gray line indicates the isotype control. **(C-H)** RNF43 hyperactiving mutants do not induce SNAP-FZD endocytosis as assessed by fluorescence microscopy. HeLa cells were co-transfected with plasmids expressing (C) SNAP-FZD5 alone, or (D) with WT RNF43 or (E-H) indicated hyperactivating mutants V162M, G166V, I186T, or R286W. SNAP-FZD was stained using SNAP-Surface549. Scale bar = 20 µm. **(I)** Hyperactivating RNF43 mutants increase the abundance of mature FZD5. Protein lysates from HEK293 cells transfected with HA-FZD5 alone or with the indicated RNF43 constructs were analysed by western blot with anti-HA antibody. Silent RNF43 mutants but not hyperactivating mutants reduce the abundance of the mature form of HA-FZD5. Tunicamycin (Tuni) was added to block N-glycosylation as indicated (Lane 3). **(J)** RNF43 forms homodimers. FLAG-tagged RNF43 was co-immunoprecipitated with Myc-tagged RNF43 in cells co-expressing both 3XFLAG-RNF43 and RNF43-MYC. **(K)** Wildtype and hyperactivating RNF43 mutants dimerize with ZNRF3. HA-ZNRF3 co-immunoprecipitated with WT or mutant Myc-FLAG-tagged RNF43. **(L)** Model: Hyperactivating mutants sequester wild-type RNF43 and/or ZNRF3.

Flow cytometry allows the assessment of membrane-bound FZD but not its intracellular fate. To further test if patient-derived RNF43 mutants are impaired in inducing FZD internalization, we utilized a SNAP-FZD5 construct (7). As reported previously, when cells expressing SNAP-FZD5 are treated with a membrane-impermeable SNAP Alexa549 reagent, the labeled FZD5 receptor is visualized at the cell surface (Figure 4C). Co-expression of RNF43 leads to rapid internalization of surface-labeled FZD5 into endocytic vesicles (Figure 4D). Supporting the flow cytometry and the Wnt/β-catenin reporter assays, co-expression of RNF43 dominant negative mutants V162M, G166V, I186T or R286W, did not lead to FZD5 endocytosis but instead enhanced its membrane localization and abundance (Figure 4E-H).

FZD proteins are glycosylated during their maturation process in the endoplasmic reticulum (ER) and Golgi. The failure of RNF43 LOF mutants to induce FZD internalization was also reflected in the relative abundance of the mature (glycosylated) and the immature (non-glycosylated) forms of FZD5 (7,33). Treatment with the glycosylation inhibitor tunicamycin collapsed the mature as well as the immature bands of FZD5 into a single band with faster mobility (Figure 4I, lane 3). Co-expression of FZD5 with wild-type RNF43 or RNF43 with silent mutations S94I, P118T or A146G led to the disappearance of the upper mature band of FZD5, indicating clearance of the mature form from the plasma membrane. In contrast, the amount of the mature FZD5 was enhanced in the presence of dominant negative RNF43 mutants (Figure 4I, lanes 8-12). The disappearance of the mature form of FZD5 in the cells expressing wild-type RNF43 or RNF43 with silent mutations S94I was rescued by treatment with bafilomycin, which blocks lysosomal degradation (Figure S4B). These results are consistent with the model that RNF43 induces FZD endocytosis and subsequent lysosomal degradation and the inactive RNF43 mutants fail to promote FZD internalization.

### The dominant negative function of RNF43 is linked to dimer formation

We explored potential mechanisms for the dominant negative, hyperactivating nature of some RNF43 mutants. Similar to our observation, three disease-associated hyperactivating RNF43 mutations have been reported previously, but the underlying mechanisms are still unclear (25). We hypothesized that RNF43 may normally form a homodimer or heterodimer with ZNRF3, and that a mutant RNF43 in the complex might inhibit the function of the wild-type endogenous protein. RNF43 dimers have not been reported previously, but its paralog ZNRF3 has been observed to form dimers in the crystal structure (34). Importantly, homodimerization is required for the activity of multiple RING domain proteins (35,36). To test whether RNF43 can form higher order complexes, we co-expressed RNF43 with two different tags, FLAG-RNF43 and RNF43-Myc. Co-immunoprecipitation experiments indeed revealed strong association of these two differently tagged RNF43 proteins, supporting the hypothesis of RNF43 homodimerization (Figure 4J). Interestingly, after removing the intracellular region of RNF43 including the RING domain, the protein could still interact with itself (Figures S4C-D). Furthermore, both WT and mutant RNF43 could interact with ZNRF3 (Figure 4K). This is consistent with a model where RNF43 normally dimerizes with itself or with ZNRF3. Dominant negative mutants may then dimerize with wild-type protein forming an inactive complex, hence reducing the clearance of FZDs from the cell surface (Figures 2 and 4) (see model Figure 4L).

### RNF43 truncation and frameshift mutations including G659fs are loss of function and predict sensitivity to PORCN inhibitor

We next examined the activity of truncation and frameshift mutations in the N-terminal half of RNF43 protein that includes the known functional domains, including the hotspot G•C track around R117. Nearly all of the N-terminal truncation mutants were either loss of function or hyperactivating (Figure 5A). Consistent with its hyperactivating activity in the Wnt/β-catenin reporter assay, RNF43 with the frameshift V287Gfs*7 in the RING domain significantly increased surface FZD abundance (Figure 5B).

**Figure 5.**
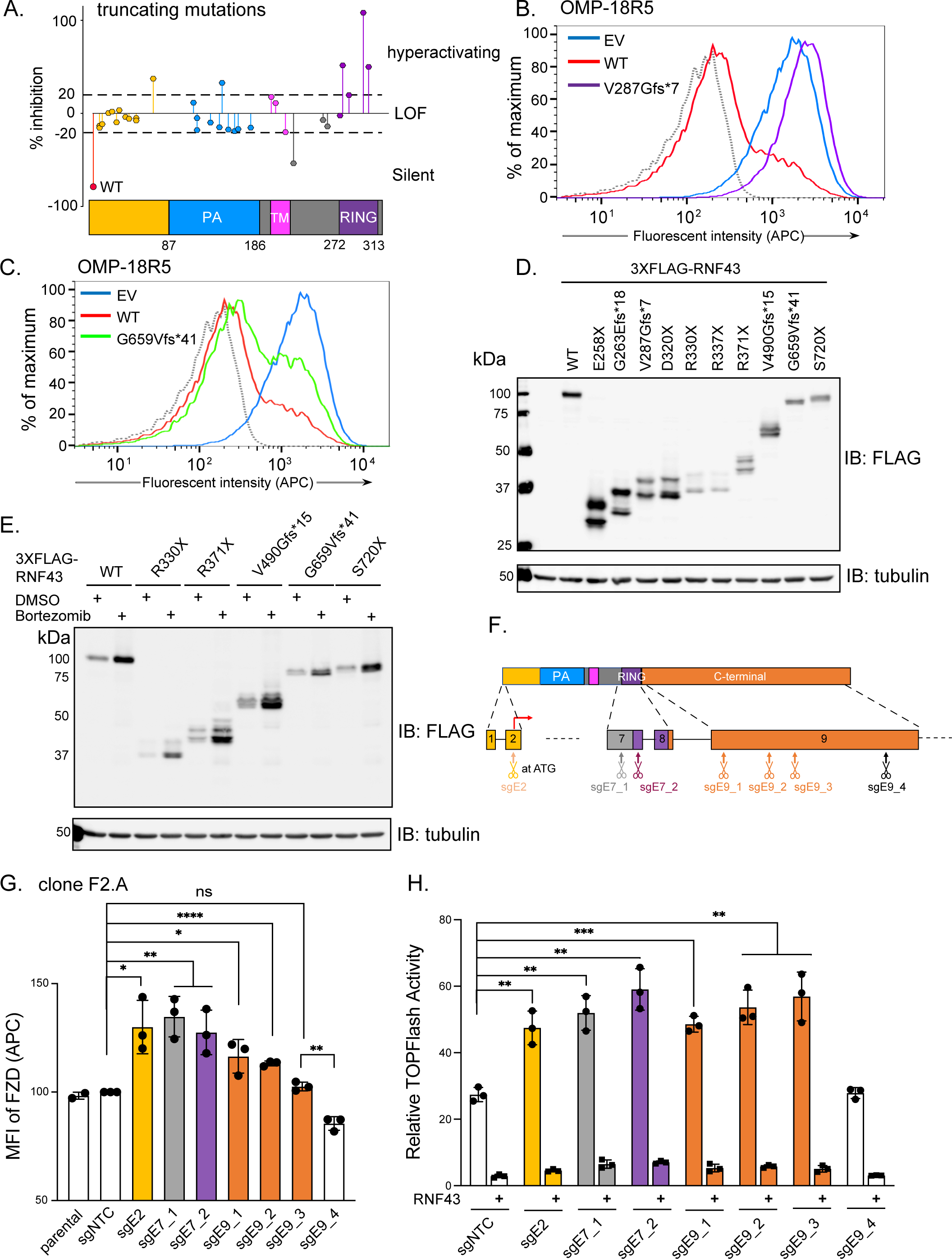
RNF43 truncating mutants are largely loss of function. **(A)** N-terminal truncation mutants in RNF43 are either loss of function or hyperactivating. Data represent percentage inhibition of WNT3A-induced Wnt/β-catenin reporter activity in HEK293 cells (as in Figure 2D) in the presence of the indicated RNF43 mutants. Data are representative of 3 independent experiments. **(B)** Expression of the hyperactivating RING domain mutant V287Gfs*7 increases cell surface FZD levels. EV, empty vector. Endogenous FZD was detected as in Figure 4A. **(C)** G659Vfs*41 RNF43 mutant is less active than WT RNF43 in regulating cell surface FZD levels. **(D)** RNF43 C-terminal truncation mutants are expressed at lower levels compared to the WT RNF43 and are regulated by proteasomal degradation. Expression of indicated 3X-FLAG RNF43 mutants in HEK293 cells was analyzed by western blotting with anti-FLAG antibody. **(E)** Selected mutants were analyzed as in D, but after transfection cells in indicated lanes were treated with proteasome inhibitor 5 nM Bortezomib overnight. **(F)** Schematic diagram of human *RNF43* genomic locus with the sgRNA targeting sites used for G-H. **(G)** CRISPR-Cas9 genome edited Panc 08.13 cell pools have increased cell surface FZD levels. Flow cytometric analysis of endogenous cell surface FZD in indicated cell pools detected using pan-FZD antibody clone F2.A. Mean fluorescence intensity (MFI) of cell pools compared to the cells transfected with non-targeting control sgRNA set to 100. Data are representative of three independent experiments. Unpaired *t-test* analysis was used for statistical analysis. **(H)** CRISPR-Cas9 genome edited Panc 08.13 cell pools have increased Wnt/β-catenin reporter activity compared to the control pools transfected with non-targeting control sgRNA. Unpaired *t-test* analysis was used for statistical analysis.

We next tested the activity of 12 C-terminal truncating mutations. In the RNF43 Wnt/β-catenin reporter assay, a number of the mutants had partial function, while others appeared to be wild-type (Figure S5A and B) as has also recently been reported (21). The G•C stretch in the C-terminus of the gene near the codon for G659 is a target for indels and along with R117fs is one of the most frequent mutations in RNF43 in multiple cancer types. Indeed, the frequency of the G659fs truncation mutant was shown to be significantly greater than predicted by chance, even in MSI cancers, suggesting they confer a growth or survival advantage to tumors harboring this mutation (5). In contrast, two recent publications concluded that indels around G659 do not compromise RNF43 activity (21,37).

To carefully examine whether G659fs and other C-terminal truncations are loss of function, we first tested the effect of G659Vfs expression on FZD cell surface abundance. Compared to the wild-type RNF43, the G659Vfs*41 mutant was modestly compromised in its ability to inhibit FZD abundance on the cell surface (Figure 5C) consistent with partial loss of function. We speculated that the C-terminal truncations and frameshifts might destabilize RNF43 protein. Since antibodies detecting endogenous RNF43 were not available, we constructed several patient-associated RNF43 truncating mutants in an N-terminal epitope-tagged construct, where a 3xFLAG was placed following the signal peptide. Indeed, we observed that while most of these constructs transcribed comparable amounts of mRNA, the mutant proteins including the recurrent G659Vfs*41 mutant were expressed at lower levels than WT RNF43 (Figure 5D). While decreased protein abundance should compromise the activity of these carboxy-terminal mutants, we were unable to detect the loss of function in the Wnt/β-catenin reporter assay (Fig S5A-B).

We speculated that *in vivo* there might be low protein abundance due to enhanced proteasomal degradation of prematurely truncated and frameshifted proteins. In fact, treating the transfected cells with the proteasome inhibitor bortezomib markedly increased the protein levels of the mutant RNF43 proteins (Figure 5E). Thus, RNF43 proteins that are prematurely truncated and/or frame-shifted may be less stable. We speculate that this difference is accentuated when these proteins are expressed endogenously.

In order to eliminate the role of exogenous overexpression, we performed genome editing of the endogenous *RNF43* gene in Panc 08.13 cells that have no underlying mutation in the Wnt/β-catenin pathway using CRISPR-Cas9. CRISPR-Cas9 mediated non-homologous end joining (NHEJ) leads to indels, resulting in frameshift mutations. The human *RNF43* gene has ten exons. Exons 7 and 8 encode the functional RING domain, whereas Exon 9 being the longest, encodes for almost half of the protein including most of the C-terminus. As illustrated in Figure 5F, we used multiple single guide RNAs (sgRNAs) to target specific regions of RNF43. We designed a sgRNA targeting the translation start site in exon 2 to serve as a positive control. To target the RING domain, we designed two sgRNAs targeting Exon 7, sgE7_1 recognizing sequences upstream of the RING domain and sgE7_2 in the RING domain. Four sgRNAs were designed to target RNF43 Exon 9 to test the impact of truncating mutations at the C-terminal of the RNF43.

Panc 08.13 cells were transfected with a plasmid expressing both Cas9 and sgRNA and selected with puromycin. The genome-editing efficiency of the sgRNAs in the cell pools was analyzed using the Surveyor nuclease assay (Figure S5D-E, Table S2). All the specific sgRNAs produced the expected bands following nuclease S treatment (arrows), indicating genome editing at the target locus. sgE9_4 sgRNA showed the same banding pattern as the non-targeting control sgRNA and turned out to be a fortuitous additional negative control, as Panc 08.13 cells were unexpectedly homozygous for a P686R polymorphism (allele frequency 0.099), altering the PAM sequence to which sgE9_4 was targeted (Figure S5F).

We then examined the consequences of targeted editing of the genomic locus of *RNF43*. Both cell surface FZD levels and endogenous Wnt/β-catenin activity were significantly increased in all the successfully edited cell pools (Figure 5G, 5H). This genome editing result stands in marked contrast to the results from transient overexpression using cDNA. We noted that *RNF43* has been reported to undergo nonsense mediated decay (NMD) in HCT116 cells (38). However, consistent with what has been reported in other systems (21) we did not observe a significant reduction in the mRNA levels in any of the successfully edited cell pools (Figure S5G) suggesting that the levels of *RNF43* in these cells are not regulated by NMD. Taken together with our prior data, we conclude that frameshift and truncation mutations in the C-terminal half of endogenous RNF43 compromise its activity, and this is in part mediated by decreasing protein stability.

### Patient derived xenografts with RNF43 C-terminal truncation mutations including P660fs have high Fzd abundance and are sensitive to PORCN inhibitor

The genome editing studies in Panc 08.13 cells demonstrated that truncating mutations in the C-terminus are loss of function, while, confusingly, transient transfection assays pointed to a different conclusion. Additional data supports the conclusion from gene editing. First, the AsPC-1 cell line with a distal C-terminal truncating mutation S720X is Wnt addicted and sensitive to PORCN inhibitors *in vivo* (19,39). To further test if chromosome-based truncating mutations in the RNF43 C-terminal region are indeed loss of function, we examined several pancreatic PDX models. We selected two models with distinct C-terminal frameshift mutations, PAXF 1861 with a heterozygous R371fs and PAXF 1869 with the recurrent P660fs mutation and loss of heterozygosity (Figure 6A). We first assessed FZD abundance by immunohistochemical (IHC) staining with a pan-FZD monoclonal antibody (Figure 6B) (39) {Pavlovic:2018ix}. Xenograft tumors from Wnt-addicted RNF43-mutant cell lines HPAF-II (E174X) and AsPC-1 (S720X) had prominent FZD staining, while two negative controls, pancreatic PDXs with wild-type RNF43, had minimal FZD staining (Fig 6B, 6C). Importantly, PAXF 1861 and PAXF 1869 both had high FZD abundance with clear membrane staining. The observation that PAXF 1861, a PDX with a heterozygous truncating mutant, had elevated cell surface FZD supports the model that inactive RNF43 mutants can also inactivate wild-type copies (Figure 4L). These data support our observations that endogenous C-terminal truncation mutations compromise the activity of RNF43 leading to increased FZD levels on the cell surface.

**Figure 6.**
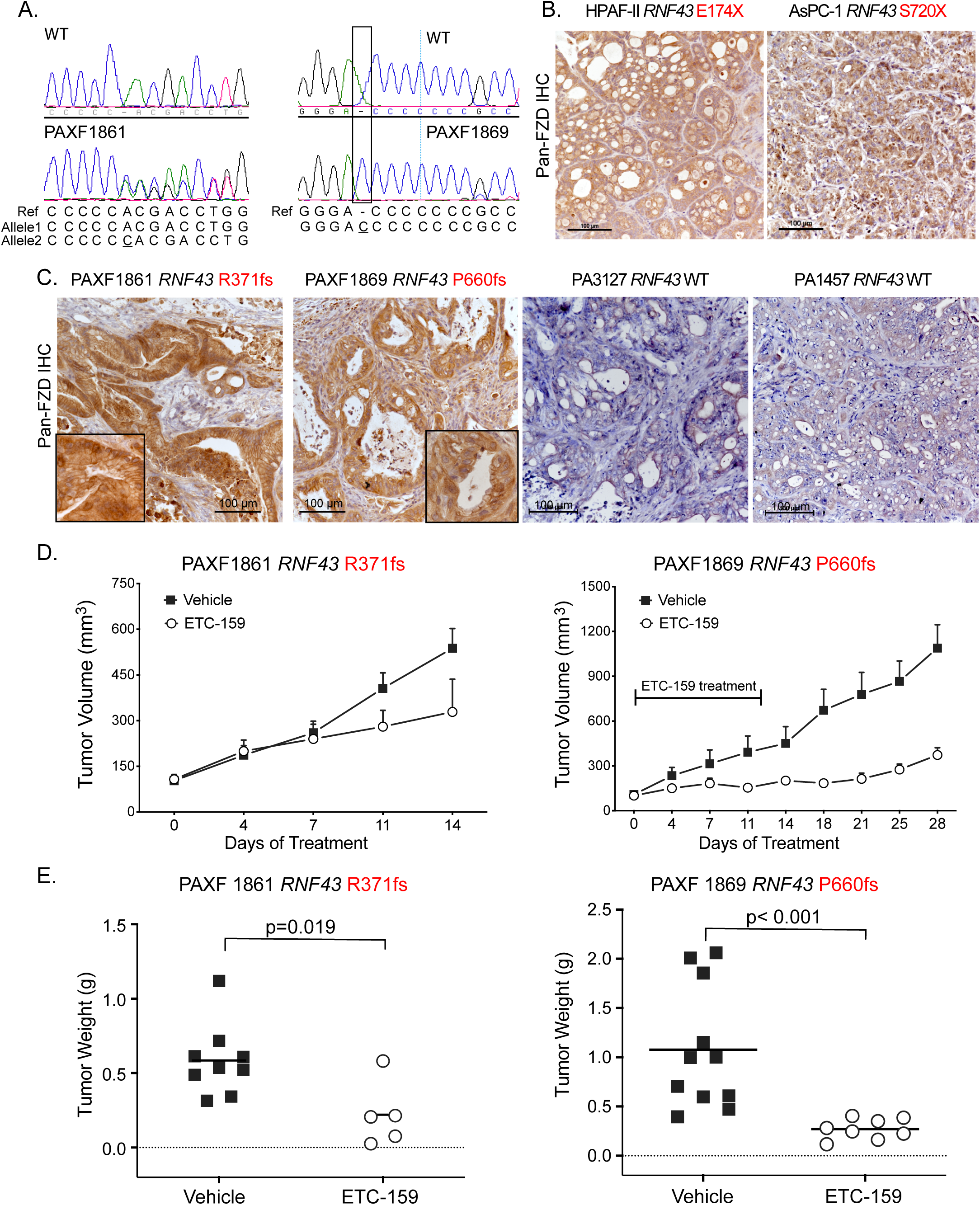
Pancreatic PDX with C-terminal truncations have increased FZD abundance and are sensitive to PORCN inhibition. **(A)** Sanger sequence traces showing the heterozygous G372fs and homozygous P660fs in the PAXF1861 and PAXF1869 pancreatic cancer patient-derived xenografts. **(B)** Increased FZD protein in HPAF-II and AsPC-1 xenografts as detected by immunohistochemical staining with pan-FZD antibody clone F2. Scale bar = 100 µm. **(C)** Pancreatic PDX PAXF1861 and PAXF1869 with RNF43 truncating mutations have high abundance of FZDs. PDX with wild-type RNF43 have barely detectable FZD protein. **(D-E)** Pancreatic PDX with C-terminal RNF43 truncating mutations are sensitive to ETC-159 treatment. NMRI-nude mice transplanted with fragments from PDX were treated with either vehicle or ETC-159. Mice implanted with PAXF 1869 were treated with ETC-159 (30 mg/kg qd) for 13 days and those implanted with PAXF 1861 were treated with 75 mg/kg qd for 15 days. Tumor weights recorded at the end of the study are shown in E. Unpaired *t-test* (Mann Whitney) was used for statistical analysis of the tumor weights.

To directly examine if these mutations are actionable, we tested the sensitivity of PDX PAXF 1869 and PAXF 1861 to the PORCN inhibitor ETC-159. The growth and final weight of tumors generated from both PAXF 1869 and PAXF 1861 in mice was significantly reduced by ETC-159 treatment (Figure 6 D-E). These results confirm that *in vivo* RNF43 C-terminal truncating mutations are loss of function.

## Discussion

RNF43 is frequently mutated in diverse cancers including colorectal, endometrial, mucinous ovarian, pancreatic and gastric (5,20,24,40,41). Wnt inhibition using upstream pathway inhibitors such as anti-FZD blocking antibodies or PORCN inhibitors is effective in preclinical cancer models with *RNF43* loss-of function mutations or R-spondin translocations, and hence these agents have advanced to clinical trials. *RNF43* is now included in sequencing panels designed to identify actionable mutations in cancer patients. However, as clinical mutations in *RNF43* span the entire length of the gene it is challenging to predict which are drivers and which are passengers. Here we analyzed a broad range of patient-derived mutations to determine those that can be used to predict Wnt addiction.

Our study consistently found that all tested nonsense and frameshift mutations, as well as a large percentage of missense *RNF43* mutations found in human cancers are either loss of function or hyperactivating, leading to increased cell surface abundance of FZDs and subsequently sensitivity to PORCN inhibition. In the absence of additional downstream Wnt pathway mutations or MSI status, these results can guide decisions as to which patients may benefit from upstream Wnt pathway inhibitors.

One unexpected finding was that a large fraction of missense and truncating mutants in the N-terminus of RNF43 acted as dominant negative mutations that hyperactivated Wnt/β-catenin signaling. These mutants failed to induce cell surface FZD receptor internalization and subsequent lysosome-mediated degradation. Mechanistically, we find that RNF43 can form a higher-order complex with itself and its paralog ZNRF3, and propose that inclusion of a mutant in the heterodimer renders the wild-type copy inactive. This suggests that cancers with one mutant *RNF43* allele, and/or tumors with mutant RNF43 that co-express ZNRF3, may still have enhanced FZD abundance at the cell surface, exhibit higher sensitivity to Wnts, and hence may benefit clinically from upstream Wnt pathway inhibitors.

We find that C-terminal truncating mutations in RNF43 also are loss of function. Importantly, we confirmed the deleterious consequences of C-terminal truncations in a MSS pancreatic cancer cell line and in several pancreatic patient-derived xenografts. We find that two xenografts with distinct C-terminal mutations P660fs and R371fs have increased cell surface FZD and are sensitive to therapeutic doses of PORCN inhibitors. This is consistent with published data that the AsPC-1 cell line with S720X is sensitive to PORCN inhibitors in xenograft models (19,39). While G659 mutations are most common in MSI cancers due to its G•C track, indels affecting the C-terminus also occur in MSS cancers and there they predict response to PORCN inhibitors. Whether RNF43 mutations predict response to PORCN inhibitors in MSI tumors with multiple additional Wnt pathway and other mutations is not known.

Why G659fs and other C-terminal truncations lose function *in vivo* but not in cell based reporter assays is not clear. The results from HEK293 cell-based reporter assays were discordant with results from both our genome editing data in Panc 08.13 cells and from functional studies in xenografts, suggesting these more complex assays may measure additional relevant properties of the C-terminus of RNF43. Our *in vitro* assays using an approach similar to others agrees with their observations (21,37). However, there are limitations of transient transfections, including the highly simplified promoter used in the *in vitro* assay, the nature of the cell types used, and that the assays are performed *in vitro* while mutant tumors grow in a far more complex environment *in vivo*. Some differences observed in the CRISPR-Cas9 edited cells may be explainable by cell type, as Li et al., used a MSI colorectal cell line KM12 with *APC* and *AXIN1* mutations. These mutations activate Wnt signaling downstream of FZD. In contrast, Panc 08.13 cells have no downstream mutations in Wnt signaling components (42). Our in-depth *in vivo* analysis of various C-terminal truncating mutations including those in the most recurrently mutated region around G659 predicts them to be loss of function and potentially actionable mutations.

Our analysis provides a comprehensive evaluation of RNF43 as a biomarker for predicting sensitivity to PORCN inhibitors. We find that all truncation or frameshift mutants are loss of function. Nearly all missense mutants in the RING domain are also loss of function. In the absence of additional downstream Wnt pathway mutations, these mutations predict actionable RNF43 mutants. The finding that simple transfection assays fail to identify a number of *in vivo* loss-of-function mutants suggests our results are actually an underestimate of the number of actionable RNF43 mutations. Increased membrane FZD abundance in *RNF43* mutant cancers has potential to identify these additional actionable tumors. Developing FZD immunohistochemical assays might further assist in selecting patients for treatment with PORCN inhibitors.

## Materials and methods

### Cell lines, plasmids and antibodies

HEK293 cells were cultured in DMEM medium supplemented with 10% fetal bovine serum, L-glutamate, pen/strep and sodium pyruvate. Panc 08.13 cells were cultured in RPMI1640 medium supplemented with 15% fetal bovine serum, L-glutamate, pen/strep, sodium pyruvate and insulin.

The HA-FZD constructs were the generous gift of Jeffrey Rubin. They contain the signal peptide of human FZD5, followed by 2XHA tags and then the corresponding mature FZD sequences. FZD1 and FZD2 are rat, FZD5 is human, and the remainder are mouse sequences. Monoclonal antibodies clone F2 and clone F2.A were obtained from Sachdev Sidhu and Stephane Angers, University of Toronto. OMP-18R5 was used as previously described (11, 43).

### Generation of RNF43 mutant constructs

A C-terminal Myc-DKK-tagged RNF43 ORF expression construct was purchased from Origene (Origene, Cat# RC214013) and verified by Sanger sequencing. Patient-derived mutations were introduced using site-directed mutagenesis following the manufacturer’s protocol (Stratagene). 3xFLAG-internal tagged RNF43 was generated by 2-step PCR, placing 3xFLAG after the signal peptide. The C-terminal Myc tagged construct used for co-immunoprecipitation studies was generated by introducing a stop codon after the sequence encoding Myc. All constructs including the patient-derived mutants were verified by Sanger sequencing.

### Wnt/β-catenin reporter assay

One day before transfection, HEK293 cells were seeded into 24-well plates coated with poly-L-lysine. Cells were transfected using Lipofectamine 2000 (Thermo Fisher) mixed with DNA in a ratio of 3:1 using the manufacturer’s protocol. For each well 100 ng TOPFlash, 50 ng mCherry, 10 ng PGK-mWNT3A and 5 ng (unless otherwise indicated) RNF43 plasmid mix was used. After 48h of transfection, cells were lysed with 100 µl Reporter Lysis Buffer (Promega) supplemented with protease inhibitor cocktail (Roche) for 20 minutes at 4°C with shaking. The cell lysates were transferred to a 96-well black plate and then an equal volume of Luciferase assay reagent (Promega) was added. Luciferase activity was measured on a Tecan Infinite 2000 plate reader. mCherry values were used for normalization.

### Immunohistochemistry staining for surface FZD

FFPE sections of the xenograft tumors were dewaxed following standard protocol. The section slides were boiled in Tris-EDTA (pH 9.0) buffer for 20 minutes for antigen retrieval. Human pan-FZD antibody clone F2 (39) {Pavlovic:2018ix} was used at 1:1000 dilution with anti-human IgG secondary antibody conjugated with HRP.

### Mouse xenograft studies

All mouse studies were approved by the institutional IACUC committee. The patient-derived xenografts studies were performed by Oncotest, Freiburg, Germany. Mouse xenograft models were established by subcutaneously injecting HPAF-II or AsPC-1 cells mixed with matrigel into NSG mice or implanting patient-derived solid tissue fragments in NMRI nude mice. ETC-159 formulated in 50% PEG400 (vol/vol) in water was administered by oral gavage at a dosing volume of 10 µL/g bodyweight. Tumors were measured by calipers and volumes calculated using the formula (length/2 × width^2). All mice were sacrificed 8 hours after the last dosing. At sacrifice, tumors were resected, weighed and snap-frozen in liquid nitrogen or fixed in 10% neutral buffered formalin.

## Acknowledgements

We gratefully acknowledge the assistance of Jamal Aliyev and other past and present members of the Virshup and Epstein labs. We thank Dr. Madelon Maurice for the SNAP-FZD5 plasmid, and Dr Jeffery Rubin for the HA-FZD constructs. We thank Dr Stephane Angers, Dr Sachdev Sidhu and University of Toronto for the pan-FZD antibodies. We acknowledge the Advanced Bioimaging facility of Singhealth/DukeNUS for assistance with the generation of imaging data. This research is supported in part by the National Research Foundation Singapore and administered by the Singapore Ministry of Health’s National Medical Research Council under the STAR Award Program MOH-000155 (to DMV). BM acknowledges the support of the Singapore Ministry of Health’s National Medical Research Council Open Fund– Independent Research grant NMRC/OFIRG/0055/2017.

## Literature Cited

1. Takada R, Satomi Y, Kurata T, Ueno N, Norioka S, Kondoh H, et al. Monounsaturated fatty acid modification of Wnt protein: its role in Wnt secretion. Dev Cell. 2006;11:791–801.

2. Coombs GS, Yu J, Canning CA, Veltri CA, Covey TM, Cheong JK, et al. WLS-dependent secretion of WNT3A requires Ser209 acylation and vacuolar acidification. J Cell Sci. 2010;123:3357–67.

3. Yu J, Chia J, Canning CA, Jones CM, Bard FA, Virshup DM. WLS retrograde transport to the endoplasmic reticulum during Wnt secretion. Dev Cell. 2014;29:277–91.

4. Sekine S, Yamashita S, Tanabe T, Hashimoto T, Yoshida H, Taniguchi H, et al. Frequent PTPRK-RSPO3 fusions and RNF43 mutations in colorectal traditional serrated adenoma. J Pathol. 2016;239:n/a–n/a.

5. Giannakis M, Hodis E, Jasmine Mu X, Yamauchi M, Rosenbluh J, Cibulskis K, et al. RNF43 is frequently mutated in colorectal and endometrial cancers. Nat Genet. 2014;46:1264–6.

6. Jiang X, Hao H-X, Growney JD, Woolfenden S, Bottiglio C, Ng N, et al. Inactivating mutations of RNF43 confer Wnt dependency in pancreatic ductal adenocarcinoma. Proc Natl Acad Sci USA. 2013;110:12649–54.

7. Koo B-K, Spit M, Jordens I, Low TY, Stange DE, van de Wetering M, et al. Tumour suppressor RNF43 is a stem-cell E3 ligase that induces endocytosis of Wnt receptors. Nature. Nature Publishing Group; 2012;488:665–9.

8. Ong CK, Subimerb C, Pairojkul C, Myint SS, Rajasegaran V, Wu Y, et al. Exome sequencing of liver fluke-associated cholangiocarcinoma. Nat Genet. 2012;44:690–3.

9. Storm EE, Durinck S, De Sousa E Melo F, Tremayne J, Kljavin N, Tan C, et al. Targeting PTPRK-RSPO3 colon tumours promotes differentiation and loss of stem-cell function. Nature. 2016;529:97–100.

10. Madan B, Virshup DM. Targeting Wnts at the source--new mechanisms, new biomarkers, new drugs. Mol Cancer Ther. 2015;14:1087–94.

11. Gurney A, Axelrod F, Bond CJ, Cain J, Chartier C, Donigan L, et al. Wnt pathway inhibition via the targeting of Frizzled receptors results in decreased growth and tumorigenicity of human tumors. Proc Natl Acad Sci USA. 2012;109:11717–22.

12. Liu J, Pan S, Hsieh MH, Ng N, Sun F, Wang T, et al. Targeting Wnt-driven cancer through the inhibition of Porcupine by LGK974. Proc Natl Acad Sci USA. 2013;110:20224–9.

13. Proffitt KD, Madan B, Ke Z, Pendharkar V, Ding L, Lee MA, et al. Pharmacological inhibition of the Wnt acyltransferase PORCN prevents growth of WNT-driven mammary cancer. Cancer Res. 2013;73:502–7.

14. Chartier C, Raval J, Axelrod F, Bond C, Cain J, Dee-Hoskins C, et al. Therapeutic Targeting of Tumor-Derived R-Spondin Attenuates β-Catenin Signaling and Tumorigenesis in Multiple Cancer Types. Cancer Res. 2016;76:713–23.

15. Hao H-X, Xie Y, Zhang Y, Charlat O, Oster E, Avello M, et al. ZNRF3 promotes Wnt receptor turnover in an R-spondin-sensitive manner. Nature. 2012;485:195–200.

16. Carmon KS, Gong X, Lin Q, Thomas A, Liu Q. R-spondins function as ligands of the orphan receptors LGR4 and LGR5 to regulate Wnt/beta-catenin signaling. Proc Natl Acad Sci USA. National Acad Sciences; 2011;108:11452–7.

17. Glinka A, Dolde C, Kirsch N, Huang Y-L, Kazanskaya O, Ingelfinger D, et al. LGR4 and LGR5 are R-spondin receptors mediating Wnt/β-catenin and Wnt/PCP signalling. EMBO Rep. John Wiley & Sons, Ltd; 2011;12:1055–61.

18. Seshagiri S, Stawiski EW, Durinck S, Modrusan Z, Storm EE, Conboy CB, et al. Recurrent R-spondin fusions in colon cancer. Nature. 2012;488:660–4.

19. Madan B, Ke Z, Harmston N, Ho SY, Frois AO, Alam J, et al. Wnt addiction of genetically defined cancers reversed by PORCN inhibition. Oncogene. 2016;35:2197–207.

20. Wang K, Yuen ST, Xu J, Lee SP, Yan HHN, Shi ST, et al. Whole-genome sequencing and comprehensive molecular profiling identify new driver mutations in gastric cancer. Nat Genet. 2014;46:573–82.

21. Li S, Lavrijsen M, Bakker A, Magierowski M, Magierowska K, Liu P, et al. Commonly observed RNF43 mutations retain functionality in attenuating Wnt/β-catenin signaling and unlikely confer Wnt-dependency onto colorectal cancers. Oncogene. 2020;485:195.

22. Bond CE, McKeone DM, Kalimutho M, Bettington ML, Pearson S-A, Dumenil TD, et al. RNF43 and ZNRF3 are commonly altered in serrated pathway colorectal tumorigenesis. Oncotarget. 2016;7:70589–600.

23. Zou Y, Wang F, Liu F-Y, Huang M-Z, Li W, Yuan X-Q, et al. RNF43 mutations are recurrent in Chinese patients with mucinous ovarian carcinoma but absent in other subtypes of ovarian cancer. Gene. 2013;531:112–6.

24. Ryland GL, Hunter SM, Doyle MA, Rowley SM, Christie M, Allan PE, et al. RNF43 is a tumour suppressor gene mutated in mucinous tumours of the ovary. J Pathol. 2013;229:469–76.

25. Tsukiyama T, Fukui A, Terai S, Fujioka Y, Shinada K, Takahashi H, et al. Molecular Role of RNF43 in Canonical and Noncanonical Wnt Signaling. Mol Cell Biol. 2015;35:2007–23.

26. Cancer Genome Atlas Research Network. Comprehensive molecular characterization of gastric adenocarcinoma. Nature. 2014;513:202–9.

27. Cerami E, Gao J, Dogrusoz U, Gross BE, Sumer SO, Aksoy BA, et al. The cBio cancer genomics portal: an open platform for exploring multidimensional cancer genomics data. Cancer Discov. American Association for Cancer Research; 2012;2:401–4.

28. Gao J, Aksoy BA, Dogrusoz U, Dresdner G, Gross B, Sumer SO, et al. Integrative Analysis of Complex Cancer Genomics and Clinical Profiles Using the cBioPortal. Sci Signal. 2013;6:pl1–pl1.

29. Chen P-H, Chen X, Lin Z, Fang D, He X. The structural basis of R-spondin recognition by LGR5 and RNF43. Genes Dev [Internet]. Cold Spring Harbor Lab; 2013;27:1345–50. Available from: http://eutils.ncbi.nlm.nih.gov/entrez/eutils/elink.fcgi?dbfrom=pubmed&id=23756651&retmode=ref&cmd=prlinks

30. Karczewski KJ, Francioli LC, Tiao G, Cummings BB, Alföldi J, Wang Q, et al. Variation across 141,456 human exomes and genomes reveals the spectrum of loss-of-function intolerance across human protein-coding genes. bioRxiv. Cold Spring Harbor Laboratory; 2019;42:531210.

31. Le DT, Uram JN, Wang H, Bartlett BR, Kemberling H, Eyring AD, et al. PD-1 Blockade in Tumors with Mismatch-Repair Deficiency. N Engl J Med. 2015;372:2509–20.

32. Mandal R, Samstein RM, Lee K-W, Havel JJ, Wang H, Krishna C, et al. Genetic diversity of tumors with mismatch repair deficiency influences anti-PD-1 immunotherapy response. Science. American Association for the Advancement of Science; 2019;364:485–91.

33. Gerlach JP, Jordens I, Tauriello DVF, van’t Land-Kuper I, Bugter JM, Noordstra I, et al. TMEM59 potentiates Wnt signaling by promoting signalosome formation. Proc Natl Acad Sci USA. National Academy of Sciences; 2018;191:201721321.

34. Zebisch M, Xu Y, Krastev C, Macdonald BT, Chen M, Gilbert RJC, et al. Structural and molecular basis of ZNRF3/RNF43 transmembrane ubiquitin ligase inhibition by the Wnt agonist R-spondin. Nat Commun. 2013;4:2787.

35. Dou H, Buetow L, Sibbet GJ, Cameron K, Huang DT. BIRC7-E2 ubiquitin conjugate structure reveals the mechanism of ubiquitin transfer by a RING dimer. Nat Struct Mol Biol. Nature Publishing Group; 2012;19:876–83.

36. Budhidarmo R, Nakatani Y, Day CL. RINGs hold the key to ubiquitin transfer. Trends Biochem Sci. 2012;37:58–65.

37. Tu J, Park S, Yu W, Zhang S, Wu L, Carmon K, et al. The most common RNF43 mutant G659Vfs*41 is fully functional in inhibiting Wnt signaling and unlikely to play a role in tumorigenesis. Sci Rep. Nature Publishing Group; 2019;9:18557–12.

38. Ivanov I, Lo KC, Hawthorn L, Cowell JK, Ionov Y. Identifying candidate colon cancer tumor suppressor genes using inhibition of nonsense-mediated mRNA decay in colon cancer cells. Oncogene. 2007;26:2873–84.

39. Steinhart Z, Pavlovic Z, Chandrashekhar M, Hart T, Wang X, Zhang X, et al. Genome-wide CRISPR screens reveal a Wnt-FZD5 signaling circuit as a druggable vulnerability of RNF43-mutant pancreatic tumors. Nat Med. 2017;23:60–8.

40. Witkiewicz AK, McMillan EA, Balaji U, Baek G, Lin W-C, Mansour J, et al. Whole-exome sequencing of pancreatic cancer defines genetic diversity and therapeutic targets. Nat Commun. 2015;6:6744.

41. Spit M, Fenderico N, Jordens I, Radaszkiewicz T, Lindeboom RGH, Bugter JM, et al. RNF43 truncating mutations mediate a tumour suppressor-to-oncogene switch to drive niche-independent self-renewal in cancer. bioRxiv. 2019;:748574.

42. Barretina J, Caponigro G, Stransky N, Venkatesan K, Margolin AA, Kim S, et al. The Cancer Cell Line Encyclopedia enables predictive modelling of anticancer drug sensitivity. Nature. 2012;483:603–7.

43. Madan B, Walker MP, Young R, Quick L, Orgel KA, Ryan M, et al. USP6 oncogene promotes Wnt signaling by deubiquitylating Frizzleds. Proc Natl Acad Sci USA. 2016;113:E2945–54.

44. Shalem O, Sanjana NE, Hartenian E, Shi X, Scott DA, Mikkelson T, et al. Genome-Scale CRISPR-Cas9 Knockout Screening in Human Cells. Science. American Association for the Advancement of Science; 2014;343:84–7.

