## supplemental information for "The functional landscape of patient derived RNF43 mutations predicts Wnt inhibitor sensitivity"

### Supporting Information

### Supplemental Methods:

##### *Analysis of mutational landscape of RNF43*

All mutations annotated as associated with RNF43 were retrieved from cBioPortal version 17.1. Mutations annotated as gene fusions or mutations annotated as affecting splicing were removed from the dataset. Truncating mutations were defined as either frameshift, or stop codon, with all others classified as missense.

To test if a specific domain was enriched or depleted for truncating or missense mutations the background mutation rate for RNF43 was calculated, i.e. the number of amino acids with a specific type of mutation divided by the length of the protein. This was then used in a Binomial test to ascertain enrichment or depletion in specific domains. All p-values were corrected using Bonferroni.

In order to identify amino acids that were mutated more frequently than expected, 1 million permutations were used to generate a null distribution to compare to the observed frequency from cBioPortal. Bonferroni correction was used to correct p-values for multiple testing.

##### *Flow cytometric analysis of endogenous FZD cell surface levels in HEK293 cells after mutant RNF43 expression.*

For detecting endogenous Frizzled levels, HEK293 or Panc 08.13 cells were seeded into 6-well plates coated with poly-L-lysine. The cells were transfected with plasmids expressing EGFP and RNF43 using TurboFect transfection reagent. After 48 h cells were stained with pan-FZD antibodies OMP-18R5 or clone F2.A. Anti-human Fc fragment APC-conjugated (Jackson lab) was used as secondary antibody.

For analyzing the expression of HA-tagged frizzleds, Panc 08.13 cells were transfected with plasmids expressing HA tagged-FZDs and RNF43 with Lipofectamine 2000. After 48 hours cells were stained with HA-tag monoclonal antibody (16B12) conjugated with Alexa Fluor 488 (A-21284, Thermo Scientific). The cells were acquired on BD LSRFortessa Cell and analyzed using FlowJo 10 software.

##### *SNAP-FZD5 cell surface labeling*

HeLa cells cultured on glass coverslips were transfected with constructs expressing SNAP-FZD5 and WT RNF43 or RNF43 mutants using lipofectamine. The cells were incubated at

37°C, 5% CO2 for 30 minutes with the labeling medium containing 5 μM SNAP-Surface 549 (NEB, S9112). Cells were then washed three times with growth medium and chased for 15 min at 37°C, 5% CO2. Cells were fixed with 4% PFA at room temperature for 15 min and mounted with medium containing DAPI . Slides were imaged with a LSM710 confocal microscope.

***Alignment of Frizzled protein sequences***

The protein sequences of ten human and mouse Frizzleds from NCBI were aligned using the web-based tool Clustal Omega (https://www.ebi.ac.uk/Tools/msa/clustalo/) from EMBL-EBL. The phylogenetic tree was generated based on the sequence alignments.

***Coimmunoprecipitation***

Transfected cells were lysed in HEPES lysis buffer (50 mM HEPES, pH 7.4, 150 mM NaCl, 0.6% IGEPAL CA-630, 1 mM EDTA, 1 mM dithiothreitol, and protease inhibitor cocktail). Cell debris was removed by centrifugation at 13,000 x *g* for 10 min at 4 ºC. 1 µg of the appropriate antibody was added to 500 µg of protein from total cell lysates and incubated at 4 ºC overnight. Subsequently, 20 µl of protein A/G plus agarose suspension beads were added for an additional 2 hours to capture the immunocomplex. For anti-FLAG immunoprecipitation, 30 µl of FLAG M2 agarose (Sigma) was used for each reaction. The pelleted immunoprecipitate was washed three times with lysis buffer, and the bound proteins were eluted with 60 µl of Laemmli buffer, boiled at 95 ºC for 5 min, and then analyzed by SDS-PAGE and immunoblotting.

***Generation of CRISPR-Cas9 edited Panc 08.13 cell pools***

Panc 08.13 cell pools with RNF43 frameshift mutations were generated via CRISPR-Cas9 genome editing. The sgRNAs targeting different RNF43 exons (primer pairs listed below) were designed using the online CRISPR design tool “GPP sgRNA Designer" from the Broad Institute (https://portals.broadinstitute.org/gpp/public/analysis-tools/sgrna-design), and cloned into pLentiCRISPRv2 system (44) . All constructs were verified by Sanger sequencing.

Panc 08.13 cells were transfected with 1 µg of pLentiCRISPRv2 with indicated respective sgRNAs using lipofectamine 2000. The transfected cells were selected for four days with 1 µg/ml of puromycin. To validate gene targeting, genomic DNA was extracted from each cell pool using QIAamp DNA Mini kit (Qiagen) and used as template to amplify targeted locus (primer pairs listed below) using Platinum Taq HiFi DNA polymerase, following the manufacturer's protocol. 15 µl of the PCR product from each gene edited cell pool was mixed with 5 µl PCR product from the cells with wild-type RNF43. The annealed PCR products were treated with CEL nuclease (Surveyor mutation detection kit, IDT, Cat #706020) The digested samples were resolved on a 10% TBE polyacrylamide gel and stained with ethidium bromide.

sgE2: Fwd: CACCGGCAGCTGGTAGCATGAGTGG
Rev: AAACCCACTCATGCTACCAGCTGCC
sgE7_1: Fwd: CACCGGCTGGCCTGGTACCTCCTGG
Rev: AAACCCAGGAGGTACCAGGCCAGCC
sgE7_2: Fwd: CACCGCAGACAGATGGCACACACAG
Rev: AAACCTGTGTGTGCCATCTGTCTGC
sgE9_1: Fwd: CACCGCGGGATGCTGGCGAATGAGG
Rev: AAACCCTCATTCGCCAGCATCCCGC
sgE9_2: Fwd: CACCGGCTATTGCACAGAACGCAGT
Rev: AAACACTGCGTTCTGTGCAATAGCC
sgE9_3: Fwd: CACCGTTAGGGCTGCAGTACACTAG
Rev: AAACCTAGTGTACTGCAGCCCTAAC
sgE9_4: Fwd: CACCGGACCAAGGATATGCCACACT
Rev: AAACAGTGTGGCATATCCTTGGTCCRNF43
RNF43 Exon 7 Fwd: GCCATGTGATCTACCTGAGTG
RNF43 Exon 7 Rev: CACTTAGCTGCATGACGTTG
RNF43 Exon 9 Fwd: GCTACAGCCATGTCTTTCTGAATGC
RNF43 Exon 9 Fwd: CTTCACCTGTCCTGAAATATTCAGCTGT

***Reverse transcription and quantitative real-time PCR***

RNA was extracted using Qiagen RNeasy mini column using the manufacturer's protocol. RNA concentration was quantified with Nanodrop2000 and 1 µg of total RNA per sample was used to make cDNA with iScript cDNA synthesis kit (BioRad). real-time PCR was performed using either ssoFast EvaGreen Supermixes (BioRad) or BlitzAmp hotstart qPCR master mix (MiRXES) on the CFX96 Real-Time PCR system (BioRad) using the following primer pairs *RNF43* (Fwd: GCTGGTGTTGCTGAAATAACTC, Rev: GAATCCAGGCTCCAGATTGTC) *HPRT1 (*Fwd: CTCCGTTATGGCGACCC, Rev: CACCCTTTCCAAATCCTCAG)

**Figure S1**

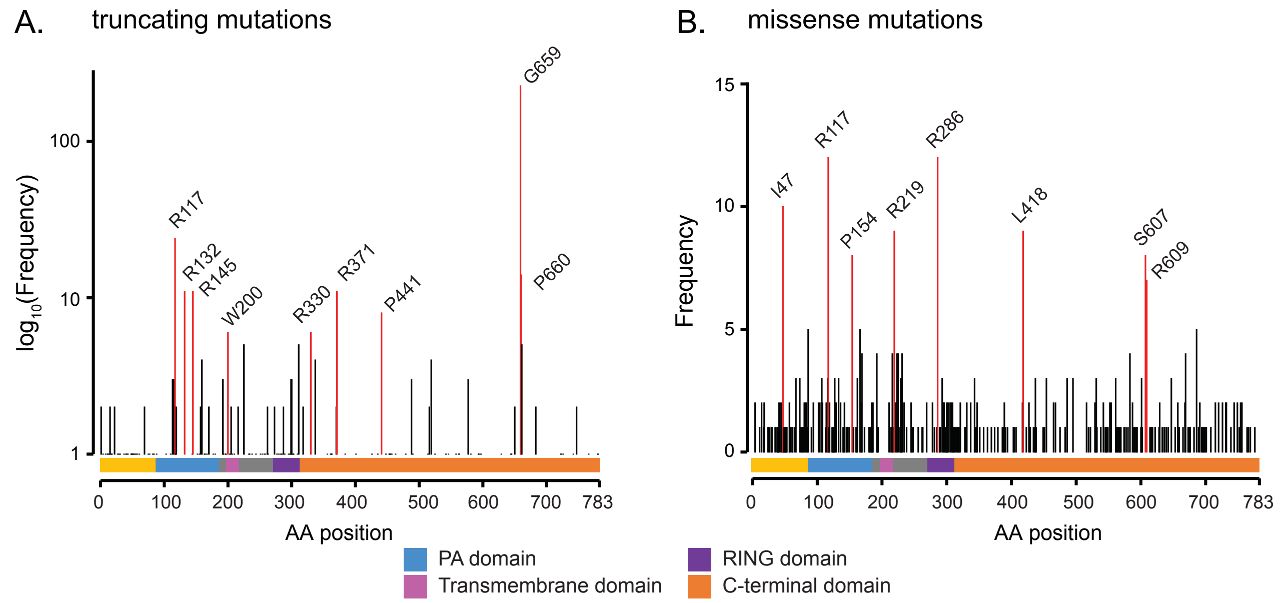
A.
truncating mutations
log10(Frequency)
100
10
1
R117
R132
R145
W200
R330
R371
P441
G659
P660
0
100
200
300
400
500
600
700
783
AA position
B.
missense mutations
Frequency
15
10
5
0
I47
R117
P154
R219
R286
L418
S607
R609
0
100
200
300
400
500
600
700
783
AA position
PA domain
Transmembrane domain
RING domain
C-terminal domain

### Supplemental Figure Legends

***Figure S1. Specific amino acids in RNF43 are recurrently mutated.***

**(A)** Permuting truncating mutations across the length of RNF43 identified several positions as been more frequently mutated than expected by chance (Bonferroni corrected p-value < 0.05 highlighted in red), including mutations in the PA domain, TM domain and at G659/P660.

**(B)** Permuting missense mutations (including common variants) over the length of RNF43 found several amino acids that were mutated more frequently than expected by chance (Bonferroni corrected p-value < 0.05, highlighted in red), including mutations in the PA and RING domain.

#### Figure S2

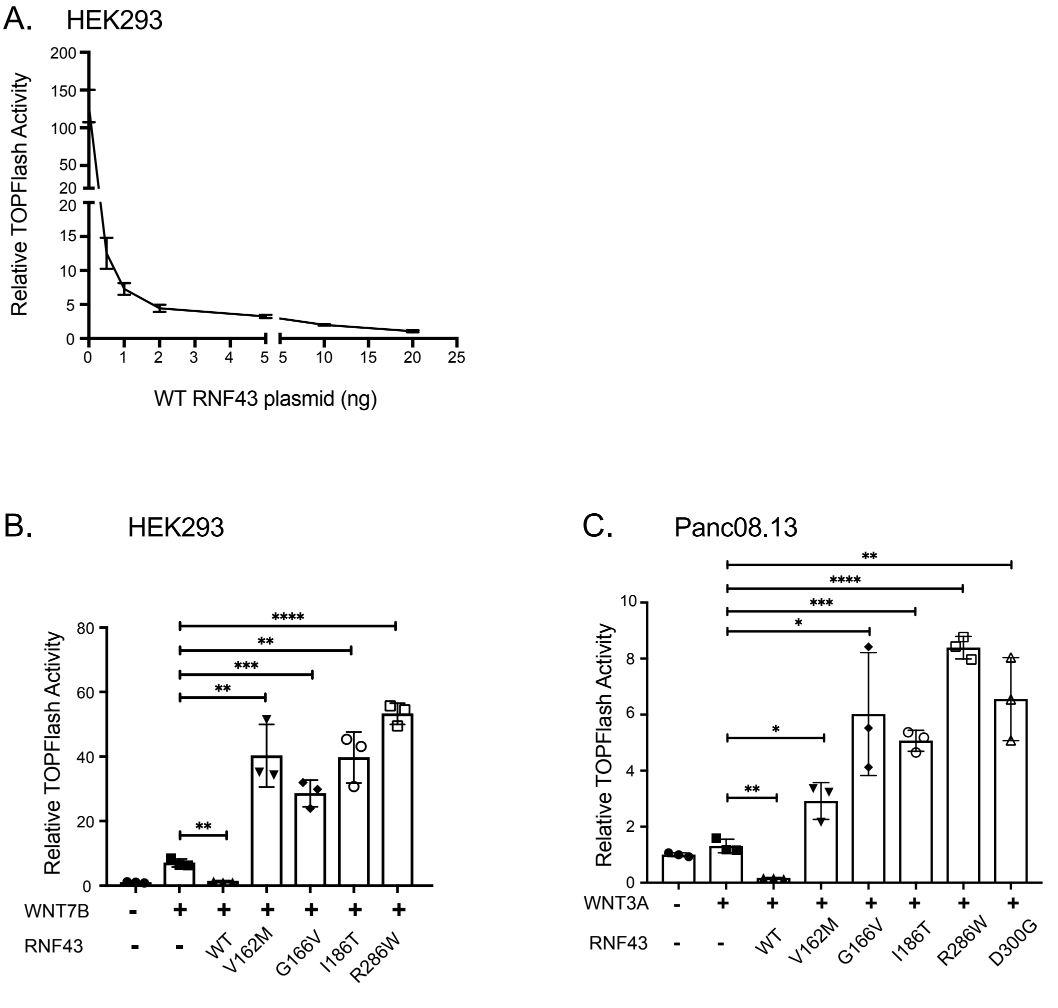

A.
HEK293
Relative TOPFlash Activity
WT RNF43 plasmid (ng)
B.
HEK293
Relative TOPFlash Activity
WNT7B
RNF43
WT
V162M
G166V
I186T
R286W
C.
Panc08.13
Relative TOPFlash Activity
WNT3A
RNF43
WT
V162M
G166V
I186T
R286W
D300G

***Figure S2. RNF43 regulates Wnt reporter activity.***

**(A)** HEK293 cells were transfected with WNT3A, Wnt/β-catenin reporter and varying concentrations of RNF43 plasmid. The relative Wnt signaling activity as measured using the luciferase assay is shown.

**(B)** Hyperactivating RNF43 mutants activate Wnt signaling induced by WNT7B. HEK293 cells were transfected with Wnt/ β-catenin reporter and plasmids expressing WNT7B and the indicated RNF43 variants. The activity of RNF43 mutants in Wnt/β-catenin reporter assay was assessed as above. Unpaired *t-test* was used for statistical analysis.

**(C)** Dominant negative RNF43 mutants activate Wnt signaling induced by WNT3A in Panc 08.13 cells. Panc 08.13 cells were transfected with Wnt/b-catenin reporter, plasmids expressing WNT3A and the indicated RNF43 variants. Unpaired *t-test* was used for statistical analysis.

#### Figure S3

```
                Intracelluar Loop 1      Intracelluar Loop 2
FZD3_HUMAN      DVTRFRYPERP 237          TITWFLAAVPKWGSEAIEKKAL  329
FZD3_MOUSE      DVTRFRYPERP 237          TITWFLAAVPKWGSEAIEKKAL  329
FZD6_HUMAN      DVRRFRYPERP 233          TITWFLAAGRKWSCEAIEQKAV  325
FZD6_MOUSE      DVRRFRYPERP 233          TITWFLAAGRKWSCEAIEQKAV  325
FZD4_HUMAN      DSSRFSYPERP 254          TLTWFLAAGLKWGHEAIEMHSS  345
FZD4_MOUSE      DSSRFSYPERP 254          TLTWFLAAGLKWGHEAIEMHSS  345
FZD9_HUMAN      EPHRFQYPERP 266          TLTWFLAAGKKWGHEAIEAHGS  356
FZD9_MOUSE      EPHRFQYPERP 267          TLTWFLAAGKKWGHEAIEAHGS  357
FZD10_HUMAN     DPARFRYPERP 262          TLTWFLAAGKKWGHEAIEANSS  352
FZD10_MOUSE     DPSRFRYPERP 263          TLTWFLAAGKKWGHEAIEANSS  353
FZD1_HUMAN      DMRRFSYPERP 354          SLTWFLAAGMKWGHEAIEANSQ  445
FZD1_MOUSE      DMRRFSYPERP 349          SLTWFLAAGMKWGHEAIEANSQ  440
FZD2_HUMAN      DMQRFRYPERP 279          SLTWFLAAGMKWGHEAIEANSQ  370
FZD2_MOUSE      DMQRFRYPERP 284          SLTWFLAAGMKWGHEAIEANSQ  375
FZD7_HUMAN      DMRRFSYPERP 288          SLTWFLAAGMKWGHEAIEANSQ  379
FZD7_MOUSE      DMRRFSYPERP 286          SLTWFLAAGMKWGHEAIEANSQ  377
FZD5_HUMAN      DMERFRYPERP 270          SLTWFLAAGMKWGNEAIAGYAQ  358
FZD5_MOUSE      DMERFRYPERP 270          SLTWFLAAGMKWGNEAIAGYAQ  358
FZD8_HUMAN      DMERFKYPERP 312          SLTWFLAAGMKWGNEAIAGYSQ  437
FZD8_MOUSE      DMERFKYPERP 309          SLTWFLAAGMKWGNEAIAGYSQ  437
                   ** *****              ::******  **.  **   .
```

```
                Intracelluar Loop 3                 proximal CTD alignment
FZD3_HUMAN      RVRIEIPLE--KENQDKLVKFMIR  420       GSKKTCFEWAS 509
FZD3_MOUSE      RVRIEIPLE--KENQDKLVKFMIR  420       GSKKTCFEWAS 509
FZD6_HUMAN      HVRQVIQHD--GRNQEKLKKFMIR  416       GSKKTCTEWAG 505
FZD6_MOUSE      HVRQVIQHD--GRNQEKLKKFMIR  416       GSKKTCTEWAG 505
FZD4_HUMAN      KIRSNLQKD--GTKTDKLERLMVK  436       WSAKTLHTWQK 506
FZD4_MOUSE      KIRSNLQKD--GTKTDKLERLMVK  436       WSAKTLHTWQK 506
FZD9_HUMAN      HIRKIMKTG--GTNTEKLEKLMVK  447       WSSKTFQTWQS 539
FZD9_MOUSE      HIRKIMKTG--GTNTEKLEKLMVK  448       WSSKTFQTWQS 540
FZD10_HUMAN     HIRRVMKTG--GENTDKLEKLMVR  443       WTSKTLQSWQQ 533
FZD10_MOUSE     HIRRVMKTG--GENTDKLEKLMVR  444       WTSKTLQSWQH 534
FZD1_HUMAN      RIRTIMKHD--GTKTEKLEKLMVR  536       WSGKTLNSWRK 632
FZD1_MOUSE      RIRTIMKHD--GTKTEKLEKLMVR  531       WSGKTLNSWRK 627
FZD2_HUMAN      RIRTIMKHD--GTKTEKLERLMVR  461       WSGKTLHSWRK 550
FZD2_MOUSE      RIRTIMKHD--GTKTEKLERLMVR  466       WSGKTLHSWRK 555
FZD7_HUMAN      RIRTIMKHD--GTKTEKLEKLMVR  470       WSGKTLQSWRR 559
FZD7_MOUSE      RIRTIMKHD--GTKTEKLEKLMVR  468       WSGKTLQSWRR 557
FZD5_HUMAN      RIRSVIKQG--GTKTDKLEKLMIR  449       WSGKTVESWRR 532
FZD5_MOUSE      RIRSVIKQG--GTKTDKLEKLMIR  449       WSGKTLESWRR 532
FZD8_HUMAN      RIRSVIKQQDGPTKTHKLEKLMIR  532       WSGKTLESWRS 615
FZD8_MOUSE      RIRSVIKQQGGPTKTHKLEKLMIR  530       WSGKTLESWRA 613
                ::*  :       : .** ::*::             : **   *
```

***Figure S3. Sequence alignment of all FZDs identifies conserved potential ubiquitylation sites.***

Frizzled intracellular loops and the proximal C-terminal domain region of all the mouse and human FZDs aligned using clustal omega identified the lysine residues that are largely conserved in all the ten Frizzleds (marked red). The less conserved lysines in the intracellular loop 3 are marked in magenta. CTD: carboxyl-terminal domain.

#### Figure S4

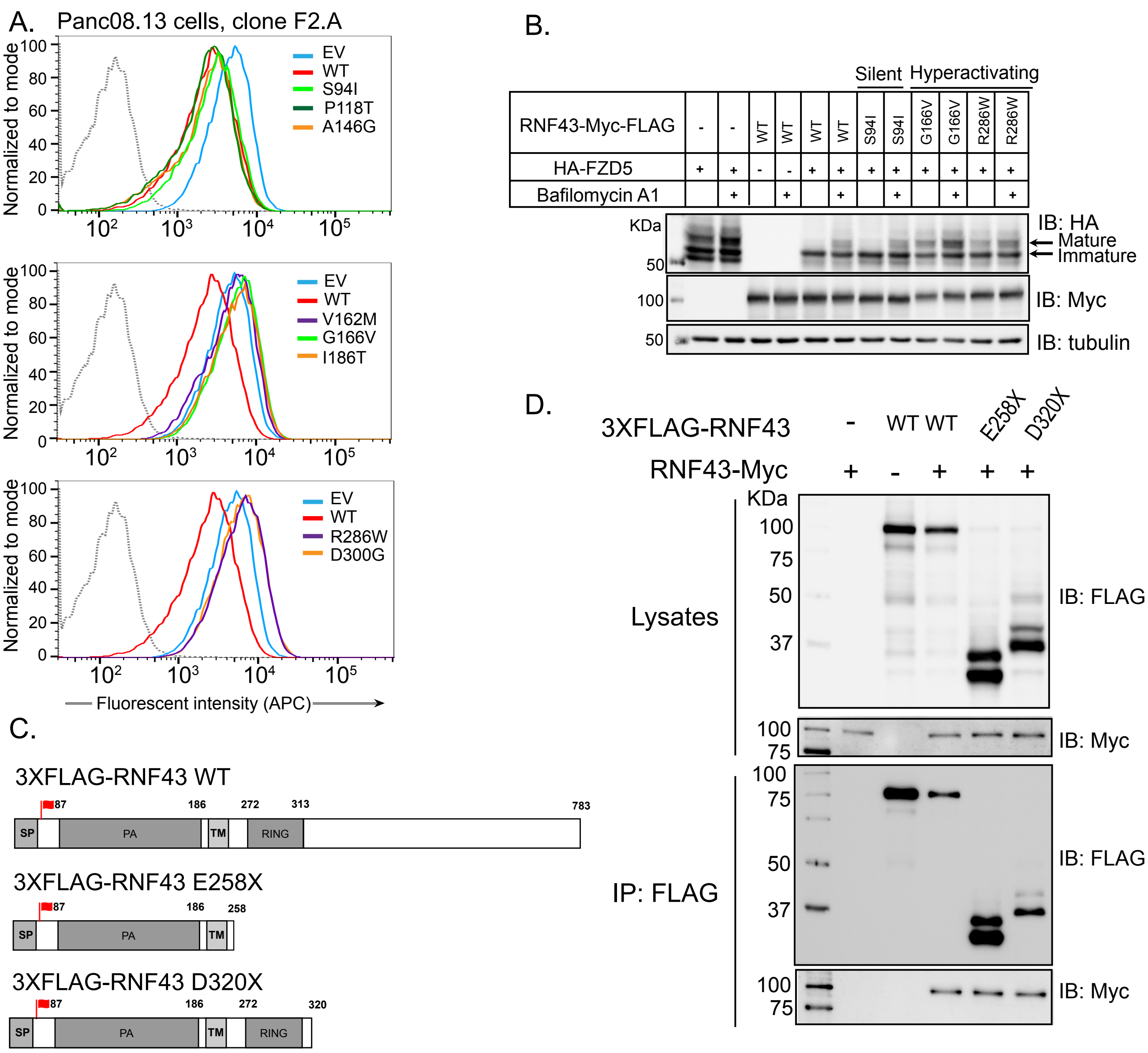

A.
Panc08.13 cells, clone F2.A
Normalized to mode
EV
WT
S94I
P118T
A146G
V162M
G166V
I186T
R286W
D300G
Fluorescent intensity (APC)
B.
Silent
Hyperactivating
RNF43-Myc-FLAG
HA-FZD5
Bafilomycin A1
KDa
IB: HA
Mature
Immature
IB: Myc
IB: tubulin
C.
3XFLAG-RNF43 WT
3XFLAG-RNF43 E258X
3XFLAG-RNF43 D320X
SP
PA
TM
RING
D.
3XFLAG-RNF43
RNF43-Myc
Lysates
IP: FLAG
IB: FLAG
IB: Myc

***Figure S4. RNF43 missense mutations regulate cell surface FZD levels.***

**(A)** Panc 08.13 were transfected with either RNF43 WT or mutant plasmids as indicated. Cell surface FZD levels were measured by staining using pan-FZD antibody clone F2.A that recognizes FZD 1, 2, 4, 5, 7 and 8. The cells co-expressing GFP were analyzed. Gray line indicates the isotype control.

**(B)** Preventing lysosomal degradation increases the accumulation of the mature form of Frizzled. HEK293 cells were transfected with HA-FZD5 with or without the indicated plasmids encoding WT or mutant RNF43. Cells were treated with 20 nM bafilomycin A1, an acidification inhibitor as indicated. Mature and immature forms of FZD5 are indicated by arrow.

**(C)** Schematic diagram showing RNF43 truncation mutants used for the assay in D.

**(D)** RNF43 self-association is independent of the carboxyl-terminal domain. Co-immunoprecipitation assays of cells co-expressing RNF43-MYC and FLAG-tagged WT RNF43 or truncation mutants.

#### Figure S5

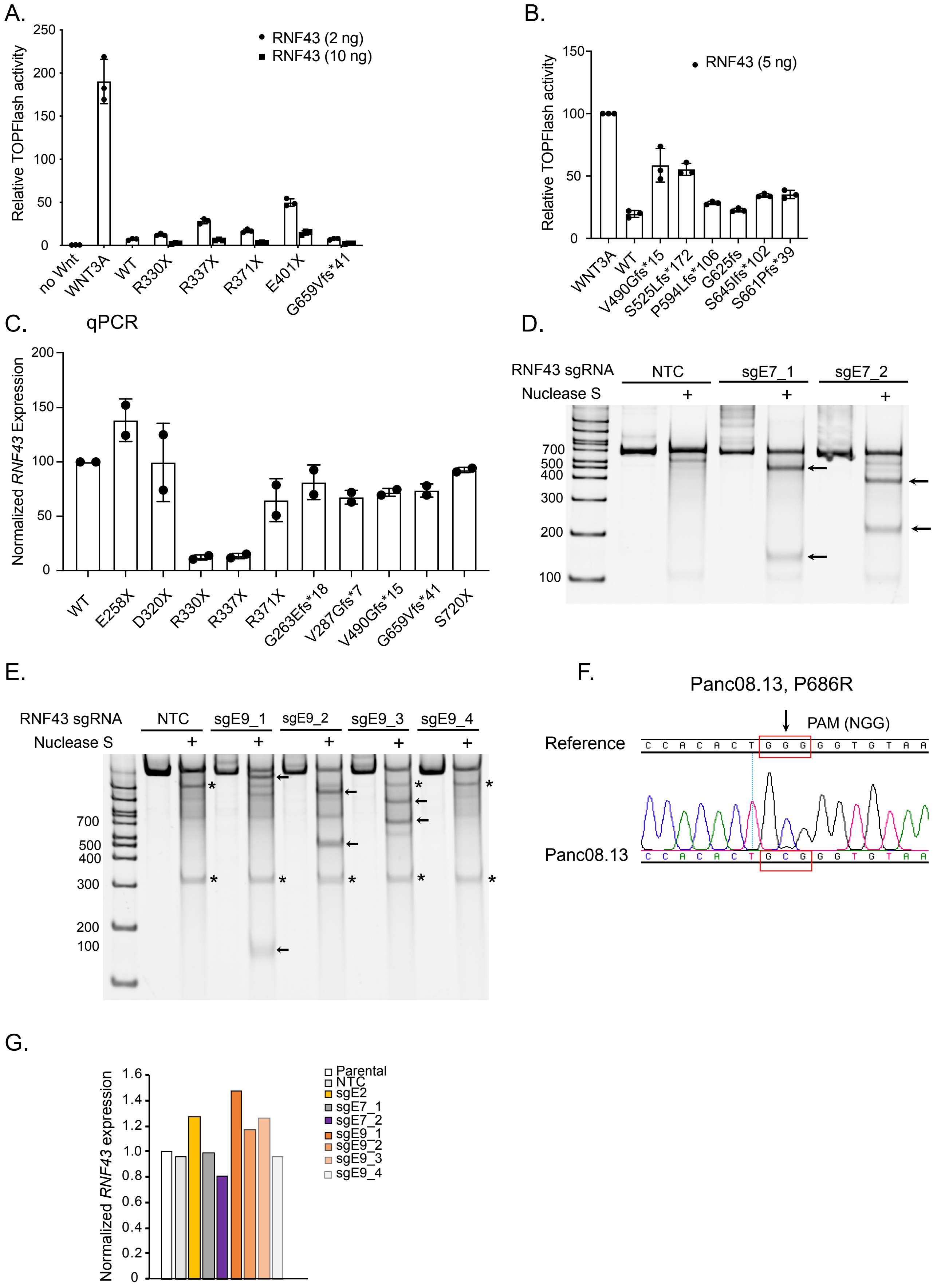
A.
Relative TOPFlash activity
250
200
150
100
50
0
RNF43 (2 ng)
RNF43 (10 ng)
no Wnt
WNT3A
WT
R330X
R337X
R371X
E401X
G659Vfs*41
B.
Relative TOPFlash activity
150
100
50
0
RNF43 (5 ng)
WNT3A
WT
V490Gfs*15
S525Lfs*172
P594Lfs*106
G625fs
S645Ifs*102
S661Pfs*39
C.
qPCR
Normalized RNF43 Expression
200
150
100
50
0
WT
E258X
D320X
R330X
R337X
R371X
G263Efs*18
V287Gfs*7
V490Gfs*15
G659Vfs*41
S720X
D.
RNF43 sgRNA
NTC
sgE7_1
sgE7_2
Nuclease S
+
700
500
400
300
200
100
E.
RNF43 sgRNA
NTC
sgE9_1
sgE9_2
sgE9_3
sgE9_4
Nuclease S
+
700
500
400
300
200
100
F.
Panc08.13, P686R
PAM (NGG)
Reference
C C A C A C T G G G G G T G T A A
Panc08.13
C C A C A C T G C G G G T G T A A
G.
Normalized RNF43 expression
1.6
1.4
1.2
1.0
0.8
0.6
0.4
0.2
0
Parental
NTC
sgE2
sgE7_1
sgE7_2
sgE9_1
sgE9_2
sgE9_3
sgE9_4

***Figure S5 C-terminal truncations are loss of function.***

**(A-B)** C-terminal RNF43 truncation mutants retain activity in *in vitro* Wnt/b-catenin reporter assay. HEK293 cells were transfected with WNT3A, Wnt/b-catenin reporter and varying concentrations of RNF43 plasmids as indicated. The relative Wnt signaling activity as measured using the luciferase assay is shown.

**(C)** mRNA levels of most of C-terminal RNF43 truncation mutants is comparable to WT.

**(D-E)** Surveyor nuclease assay to assess Cas9-sgRNA cleavage. Samples were resolved on a 10% PAGE gel containing ethidium bromide. Arrows indicate the correct fragments.

* indicates potential polymorphism.

**(F)** Sanger sequence traces showing the P686R polymorphism in the Panc08.13 cells. PAM sequence (GGG) is highlighted by the red box.

**(G)** RNF43 mRNA does not undergo nonsense mediated decay in the Panc 08.13 CRISPR-Cas9 edited Panc 08.13 cell pools.

**Table S1: A summary of 130 disease-associated RNF43 mutants profiled in the current study.**

The results are presented as percentage inhibition (% inhibition). The Wnt/β-catenin reporter activity in the presence of WNT3A alone was set to baseline. Wild-type RNF43 produces 80%-90% inhibition. Loss of function mutants were defined as producing only -20% to 20% inhibition. Partial Loss-of-function mutants have -50% to -20% inhibition in signaling. Mutants with activity 20% above baseline were classified as hyperactivating.

**Missense Mutants**

| AA change | AA position | Tumor Type | Percentage inhibition | Activity |
|---|---|---|---|---|
| A11S | 11 | Pancreatic adenocarcinoma | -75.6 | WT |
| P14S | 14 | Pancreatic adenocarcinoma | -71.0 | WT |
| L17M | 17 | Pancreatic adenocarcinoma | -82.9 | WT |
| M18V | 18 | Clear cell renal cell carcinoma | -84.7 | WT |
| S41T | 41 | Bladder Urothelial Carcinoma | -18.0 | LOF |
| I48V | 48 | Oesophagus squamous cell carcinoma, ovarian | -72.3 | WT |
| V50A | 50 | Colorectal Adenocarcinoma | -56.1 | WT |
| L61M | 61 | Breast Invasive Ductal Carcinoma | -86.3 | WT |
| G67C | 67 | Colorectal Adenocarcinoma | 104 | hyperactivating |
| G67D | 67 | Colorectal Adenocarcinoma | 139 | hyperactivating |
| F69C | 69 | Pancreatic (cell line) | 34 | hyperactivating |
| G80E | 80 | Glioblastoma Multiforme | 89 | hyperactivating |
| G80R | 80 | Ampullary Carcinoma | 123 | hyperactivating |
| L82S | 82 | Stomach Adenocarcinoma, Rectal Adenocarcinoma | -17.8 | LOF |
| M83T | 83 | Colorectal Adenocarcinoma | -83.6 | WT |
| S85F | 85 | uterine Endometrioid Carcinoma | -74.8 | WT |
| S94I | 94 | Pancreatic adenocarcinoma | -94.2 | WT |
| G102E | 102 | Leukaemia | -74.6 | WT |
| V107I | 107 | Lung Adenocarcinoma | -80.2 | WT |
| V107L | 107 | hairy cell leukaemia | -70.8 | WT |
| R113L | 113 | Lung Adenocarcinoma | -71.6 | WT |
| R117S | 117 | Hepatocellular Carcinoma | -78.1 | WT |
| P118T | 118 | Ovarian cancer | -91.2 | WT |
| C119Y | 119 | Ampullary Carcinoma | 16.1 | LOF |
| R127P | 127 | Pancreatic adenocarcinoma | 108.5 | hyperactivating |
| R127Q | 127 | Pancreatic adenocarcinoma | -87.5 | WT |
| M128V | 128 | patient | -78.8 | WT |
| G133E | 133 | Colorectal Adenocarcinoma | 75.8 | hyperactivating |
| G133R | 133 | Pancreatic adenocarcinoma | -46.7 | partial LOF |
| A146G | 146 | Ovarian cancer | -90.5 | WT |

| | | | | |
|---|---|---|---|---|
| P154L | 154 | Pancreatic adenocarcinoma, Colorectal Adenocarcinoma, endometrial | 54.3 | hyperactivating |
| V162M | 162 | Pancreatic adenocarcinoma, colorectal adenocarcinoma | 52 | hyperactivating |
| G166V | 166 | Pancreatic adenocarcinoma | 97 | hyperactivating |
| A169T | 169 | Colorectal Adenocarcinoma, intraductal Papillary Mucinous Neoplasm, Urethral Adenocarcinoma, pancreatic | 19.0 | LOF |
| E170K | 170 | Colorectal Adenocarcinoma | -92.9 | WT |
| K171R | 171 | Oesophagus squamous cell carcinoma | -12.4 | LOF |
| M173T | 173 | Renal Clear Cell Carcinoma | -84.8 | WT |
| H183R | 183 | Endometrium | -90.3 | WT |
| R185M | 185 | Autonomic ganglia Neuroblastoma | -85.2 | WT |
| I186T | 186 | Pancreatic adenocarcinoma | 144 | hyperactivating |
| V211A | 211 | small cell lung cancer | -83.5 | WT |
| S216L | 216 | Colorectal Adenocarcinoma | -91.1 | WT |
| V217M | 217 | Pancreatic adenocarcinoma | -59.0 | WT |
| R219H | 219 | stomach Adenocarcinoma, colorectal adenocarcinoma, uterine endometrioid carcinoma | -87.5 | WT |
| R221Q | 221 | Colorectal Adenocarcinoma | -81.5 | WT |
| R221W | 221 | Colorectal Adenocarcinoma | -74.6 | WT |
| R223C | 223 | Colorectal Adenocarcinoma, cutaneous melanoma | -82.3 | WT |
| R225H | 225 | glioblastoma multiforme, Colorectal Adenocarcinoma | -82.3 | WT |
| S227R | 227 | Liver Neoplasm | -87.0 | WT |
| R286Q | 286 | stomach Adenocarcinoma, colorectal adenocarcinoma, pancreatic adenocarcinoma, | 91.3 | hyperactivating |
| R286W | 286 | Pancreatic Adenocarcinoma, Colorectal Adenocarcinoma, endometrial | 140 | hyperactivating |
| C298Y | 298 | Hepatocellular carcinoma | 170 | hyperactivating |
| D300G | 300 | Stomach Adenocarcinoma | 74 | hyperactivating |
| D300Y | 300 | Hepatocellular carcinoma, lung non-small cell carcinoma | 194 | hyperactivating |
| P301S | 301 | Head & Neck Squamous cell carcinoma | 170 | hyperactivating |
| Q305R | 305 | Stomach Adenocarcinoma | -85.0 | WT |
| R307W | 307 | Endometrium | 125 | hyperactivating |
| C309F | 309 | Pancreatic adenocarcinoma, medulloblastoma | 109 | hyperactivating |
| P310A | 310 | Pancreatic adenocarcinoma | 135 | hyperactivating |
| E318D | 318 | Desmoplastic Melanoma | -86.5 | WT |
| G319R | 319 | melanoma | -78.0 | WT |
| P328S | 328 | Colorectal Adenocarcinoma | -89.3 | WT |
| Q344H | 344 | Hepatocellular Carcinoma | -74.6 | WT |

| | | | | |
|---|---|---|---|---|
| Y357C | 357 | Uterine Endometrioid Carcinoma | -88.5 | WT |
| R389C | 389 | Colorectal Adenocarcinoma, lymphoma | -74.8 | WT |
| R389H | 389 | Bladder Urothelial Carcinoma, colorectal adenocarcinoma | -60.9 | WT |
| P391S | 391 | Cutaneous melanoma | -90.3 | WT |
| G447E | 447 | Colorectal Adenocarcinoma | -98.2 | WT |
| R454C | 454 | Colorectal Adenocarcinoma | -76.2 | WT |
| D465N | 465 | Colorectal Adenocarcinoma | -48.9 | partial LOF |
| C471Y | 471 | Oesophagus Adenocarcinoma | -82.0 | WT |
| S474P | 474 | Pancreatic ductal adenocarcinoma | -23.2 | partial LOF |
| V479L | 479 | Bladder Cancer | -84.3 | WT |
| T483M | 483 | Colorectal Adenocarcinoma, Nasopharyngeal carcinoma | -32.2 | partial LOF |
| S486C | 486 | Stomach Adenocarcinoma | -49.5 | partial LOF |
| S486I | 486 | Pancreatic adenocarcinoma | -60.0 | WT |
| S495Y | 495 | Colorectal Adenocarcinoma | -61.1 | WT |
| D516G | 516 | Stomach Adenocarcinoma | -91.1 | WT |
| R531C | 531 | Colorectal Adenocarcinoma, Cutaneous Melanoma | -68.8 | WT |
| R531H | 531 | Endometrium, | -77.6 | WT |
| R567K | 567 | Colorectal Adenocarcinoma | -67.3 | WT |
| S593F | 593 | Pancreatic Adenocarcinoma | -78.4 | WT |
| S607L | 607 | Breast Invasive Carcinoma, lung squamous cell carcinoma, uterine endometroid carcinoma, bladder urothelial carcinoma | -83.8 | WT |
| S720L | 720 | Pancreatic adenocarcinoma | -93.6 | WT |
| R755S | 755 | Pancreatic adenocarcinoma | -90.3 | WT |

#### Truncation Mutants

| AA change | AA position | Tumor Type | Percentage inhibition | Activity |
|---|---|---|---|---|
| Q8X | 8 | Pancreatic adenocarcinoma | -13.3 | LOF |
| Q8Afs*24 | 8 | Pancreatic adenocarcinoma | -15.5 | LOF |
| A11Lfs*27 | 11 | Pancreatic adenocarcinoma | -11.5 | LOF |
| M18Ifs*23 | 18 | Pancreatic (cell line) | 0.4 | LOF |
| Q22X | 22 | Pancreatic adenocarcinoma | 2 | LOF |
| R27Dfs*25 | 27 | Colorectal Adenocarcinoma | -9.6 | LOF |
| L30fs*20 | 30 | patient | 3.4 | LOF |
| E37X | 37 | Colorectal Adenocarcinoma | -3.8 | LOF |
| S41X | 41 | Mucinous Cystic Neoplasm | -5.7 | LOF |
| R49Sfs*25 | 49 | Stomach Adenocarcinoma | -5.1 | LOF |
| R49fs | 49 | Pancreatic ductal adenocarcinoma PDX | -7.7 | LOF |
| V68fs | 68 | Pancreatic ductal adenocarcinoma PDX | 37.3 | hyperactivating |

| | | | | |
|---|---|---|---|---|
| R113X | 113 | Colorectal Adenocarcinoma, intraductal Papillary Mucinous Neoplasm | 11.5 | LOF |
| R117Afs*41 | 117 | stomach Adenocarcinoma, colorectal adenocarcinoma, ampullary carcinoma | -5.0 | LOF |
| R117Pfs*8 | 117 | stomach Adenocarcinoma, colorectal adenocarcinoma | -18.0 | LOF |
| R132X | 132 | Pancreatic, Colorectal Adenocarcinoma, Gastric | -9.9 | LOF |
| T142fs | 142 | Pancreatic ductal adenocarcinoma PDX | -15.0 | LOF |
| R145X | 145 | Pancreatic adenocarcinoma, Colorectal Adenocarcinoma, esophageal adenocarcinoma, stomach adenocarcinoma, bladder Adenocarcinoma, Uterine Endometrioid Carcinoma | 32.8 | hyperactivating |
| Q152X | 152 | Intraductal Papillary Mucinous Neoplasm | -17.4 | LOF |
| W159X | 159 | Pancreatic adenocarcinoma, Colorectal Adenocarcinoma, Mucinous Stomach Adenocarcinoma | -19.1 | LOF |
| L163Ffs*5 | 163 | Pancreatic adenocarcinoma | -16.8 | LOF |
| Y177X | 177 | Intraductal Papillary Mucinous Neoplasm | -15.3 | LOF |
| W200X | 200 | Colorectal Adenocarcinoma, cutaneous melanoma, glioblastoma multiforme | 17.8 | LOF |
| V205Wfs*7 | 205 | Pancreatic adenocarcinoma | 11.1 | LOF |
| S216X | 216 | Intraductal Papillary Mucinous Neoplasm, Prostate Adenocarcinoma | -20.1 | LOF |
| R225Pfs*34 | 225 | Colorectal Adenocarcinoma | -53.6 | WT |
| E258X | 258 | Stomach Adenocarcinoma | -7.1 | LOF |
| G263Efs*18 | 263 | Pancreatic Adenocarcinoma, Colorectal Adenocarcinoma | -14.0 | LOF |
| E277X | 277 | Stomach Adenocarcinoma | -2.3 | LOF |
| S280X | 280 | Colorectal Adenocarcinoma | 51.7 | hyperactivating |
| V287Gfs*7 | 287 | Stomach Adenocarcinoma | 19.7 | LOF |
| L303Ffs*140 | 303 | Pancreatic adenocarcinoma | 108.2 | hyperactivating |
| C309Afs*110 | 309 | Pancreatic adenocarcinoma | 50 | hyperactivating |
| R330X | 330 | Pancreatic, Colorectal Adenocarcinoma, Gastric | -81.7 | WT |
| R337X | 337 | Ovarian, Endometrium, Oesophagus | -55.5 | WT |
| R371X | 371 | Pancreatic, Colorectal Adenocarcinoma | -84.1 | WT |
| E401X | 401 | Stomach Adenocarcinoma | -39.3 | partial LOF |
| V490Gfs*15 | 490 | Pancreatic adenocarcinoma | -36.7 | partial LOF |
| S525Lfs*172 | 525 | Colorectal Adenocarcinoma | -44.7 | partial LOF |
| P594Lfs*106 | 594 | Ampullary Carcinoma | -93.0 | WT |
| G625fs | 625 | Pancreatic ductal adenocarcinoma PDX | -77.3 | WT |
| S645Ifs*102 | 645 | Esophageal Adenocarcinoma | -66.4 | WT |

| | | | | |
|---|---|---|---|---|
| G659Vfs*41 | 659 | Pancreatic, Colorectal Adenocarcinoma, Esophageal Adenocarcinoma, Lung Squamous Cell Carcinoma, Ampullary Carcinoma, stomach adenocarcinoma, Prostate Adenocarcinoma, Uterine Endometrioid Carcinoma, Breast Invasive Ductal Carcinoma, Bladder Urothelial Carcinoma | -94.9 | WT |
| S661Pfs*39 | 661 | Stomach Adenocarcinoma, Acinar Cell Carcinoma of the Pancreas | -95.0 | WT |
| S720X | 720 | Pancreatic (cell line) | -91.4 | WT |

#### Table S2: Surveyor nuclease assay expected cleavage products

| Primer pair | Amplicon length (bp) | sgRNA_1 expected fragment size (bp) | gRNA_2 expected fragment size (bp) | gRNA_3 expected fragment size (bp) | gRNA_4 expected fragment size (bp) |
|---|---|---|---|---|---|
| RNF43 exon7 | 626 | 154, 472 | 229, 397 | NA | NA |
| RNF43 exon9 | 1506 | 149, 1357 | 489, 1017 | 651, 855 | 1188, 318 |
